# Quantitative Motion-Corrected PALM Links Endosome Structure and Dynamics in Live Cells

**DOI:** 10.64898/2026.06.28.735082

**Authors:** Yu Xu, Santosh Adhikari, Elias M. Puchner

## Abstract

Quantitative structural analysis by Photoactivated Localization Microscopy (PALM) on the nanoscale is often restricted to fixed cells because motion during prolonged data acquisition distorts image reconstruction. Here, we develop motion-corrected PALM (mcPALM), a live-cell super-resolution approach combining a conventional fluorescence channel with PALM to correct motion-induced spreading of localizations. We further introduce a photoactivation-based correction to estimate molecule numbers from incomplete trajectories. Using PI3P-marked endosomes in yeast as a dynamic model system, we show that mcPALM recovers a live-cell maturation trajectory linking motion-corrected endosome size and calibrated PI3P content, consistent with fixed-cell benchmarks. Unlike fixed-cell PALM, mcPALM preserves endosome dynamics, revealing stage-dependent directed transport and maturation-associated motility shift. Thus, mcPALM extends PALM from static structural measurements in fixed samples to integrated quantification of nanoscale structure, molecular composition and dynamics in living cells. This framework is broadly applicable to other mobile organelles and biomolecular assemblies, enabling live-cell studies on how molecular organization and dynamics are coupled to biological function.

## Introduction

Many intracellular structures that control cell physiology are both small and dynamic. Endocytic compartments (Puchner et al., 2013; Rink et al., 2005; Toshima et al., 2006), lipid droplets (Adhikari et al., 2019, 2021), signaling clusters (Belyy et al., 2020; Calebiro et al., 2013), and other molecular assemblies often have nanoscale dimensions while undergoing rapid motion and dynamic remodeling in living cells. A full understanding of these systems therefore needs more than a static measurement of position or intensity. Instead, it requires linking morphology, molecular content, and dynamic behavior of the same structure over time. However, integrating these measurements in living cells remains challenging.

Conventional fluorescence microscopy is well suited for live-cell imaging and has been widely used to monitor the spatial distribution, motion, and interaction of intracellular organelles. However, many intracellular structures, such as endocytic vesicles and endosomes, are smaller than or comparable to the optical diffraction limit of ∼200 nm, limiting the ability of conventional imaging to resolve their detailed morphology or to quantify their molecular content (Prescianotto-Baschong and Riezman, 1998; Huang et al., 2009). Single-molecule localization microscopy methods, including Photoactivated Localization Microscopy (PALM), overcome this limitation by sparsely activating photoactivatable fluorescent proteins (PAFPs) to avoid their spatial overlap, precisely localizing their centers, and accumulating many localization events into a super-resolution reconstruction with ∼20 nm localization precision (Betzig et al., 2006; Hess et al., 2006; Rust et al., 2006; Thompson et al., 2002; Mortensen et al., 2010). This 10-fold improvement enables structural analysis below the optical diffraction limit and, with appropriate correction, single-molecule counting (Lee et al., 2012; Puchner et al., 2013). Nevertheless, PALM typically requires data acquisition over thousands of frames. In live cells, this spreads out single molecule localizations along the trajectory of moving structures, producing distorted reconstructions as well as unreliable size and molecule number measurements. For this reason, PALM is often performed in chemically fixed cells to preserve reconstruction quality. Fixation, however, eliminates the dynamic information that is central to understanding many intracellular processes.

The intracellular endocytic transport system provides a particularly useful biological context in which to address this limitation. After internalization from the plasma membrane, cargo molecules are transported through vesicular and endosomal compartments whose lipid and protein compositions are dynamically regulated. Phosphoinositides are critical determinants of membrane identity, and phosphatidylinositol 3-phosphate (PI3P), as one of them, is a key regulator of endosome maturation (Posor et al., 2022; Di Paolo and De Camilli, 2006; He et al., 2017; Marat and Haucke, 2016). PI3P recruits effector proteins containing FYVE or PX domains (Simonsen et al., 1998; Cabrera et al., 2014; Nielsen et al., 2000) and interacts with Rab GTPases, which together coordinate tethering, fusion, cargo sorting, and progression from early to late endosomal states (Rink et al., 2005; Langemeyer et al., 2018; Christoforidis et al., 1999; Tremel et al., 2021; Simonsen et al., 1998; Orr and Wickner, 2023; Balderhaar and Ungermann, 2013; Lachmann et al., 2011). Thus, endosome maturation is a multidimensional process involving coordinated changes in molecular composition, size, morphology, motion, and membrane remodeling.

Previous live-cell and fixed-cell imaging studies have provided complementary but incomplete views of this process. Conventional fluorescence microscopy can capture time-dependent movement, membrane remodeling, and transition or progression of endosomes in yeast (Day et al., 2018; Arlt et al., 2015; Toshima et al., 2006, 2016; Miyashita et al., 2018; Casler and Glick, 2020) and mammalian cells (He et al., 2017; Rink et al., 2005; Zajac et al., 2013), but it lacks the spatial resolution needed to resolve morphology and to measure absolute molecule number. Fixed-cell PALM, on the other hand, can quantify endosome size and molecule content at the single molecule level (Puchner et al., 2013), but it cannot connect these structural and molecular features to dynamic behaviors of endosomes. As endosome dynamics such as actin-dependent transport modes and transitions across motion states are emerging as important metrics for endosome identity and maturation (Toshima et al., 2006, 2016), a key challenge remains to integrate nanoscale structure, molecular counting, and dynamics within individual moving endosomes in live cells.

Here, we develop live-cell motion-corrected PALM (mcPALM) to address this challenge. mcPALM combines a conventional fluorescence reference channel with a PALM localization channel, allowing the trajectory of a moving target to be subtracted from its single molecule localization coordinates and thereby correcting motion-induced blurring of the super-resolution image. In this study, we apply mcPALM to PI3P-marked endosomes in living yeast cells as a model system and extend mcPALM into a quantitative framework for simultaneously extracting key maturation-related features, including nanoscale size, PI3P content, mobility states, membrane remodeling events, and Rab-defined maturation stages, from individual moving endosomes. We show that mcPALM recovers compact endosome morphologies from motion-induced distortions and measures accessible PI3P content using a correction for single molecule counting based on the photoactivation energy. These live-cell measurements reproduce the two-phases of the endosome maturation trajectory previously obtained only from fixed cells, showing that PI3P accumulation precedes endosome enlargement. Unlike fixed-cell PALM, mcPALM further connects this structural and molecular trajectory with live-cell dynamic information, revealing stage-dependent directed transport modes, maturation-associated motility shifts, fusion-associated remodeling events, and distinct stages marked by Rab GTPases. Together, this work establishes mcPALM as a quantitative live-cell super-resolution imaging strategy for linking nanoscale structure, molecular composition, and dynamics of moving intracellular objects and for fully decoding intrinsically dynamic intracellular processes.

## Results

### mcPALM corrects distorted PALM images of moving endosomes in living cells

To establish mcPALM, we chose endosomal PI3P as our target, which is a defining lipid on endosomal membranes and serves as a key regulator for protein recruitment and endosome maturation (Posor et al., 2022; Marat and Haucke, 2016; Cabrera et al., 2014; Nielsen et al., 2000; Orr and Wickner, 2023). We tagged a tandem version of the EEA1-derived FYVE domain with both GFP and mEos2 (Puchner et al., 2013; Patki et al., 1998; Gillooly, 2000), which specifically binds to exposed PI3P sites on endosomal membranes and can perform stable detection of accessible PI3P pools (Fig. 1A). As in previous work (Puchner et al., 2013), FYVE-mEos2 and FYVE-GFP were expressed from the weak pCyc promoter to minimize possible perturbation of the endocytic pathway. This dual-channel design allowed GFP to report the diffraction-limited position and motion of PI3P-positive endosomes, and allowed mEos2 to provide sparse single molecule localizations for the PALM reconstruction of endosome morphologies (Fig. 1A).

**Figure 1.**
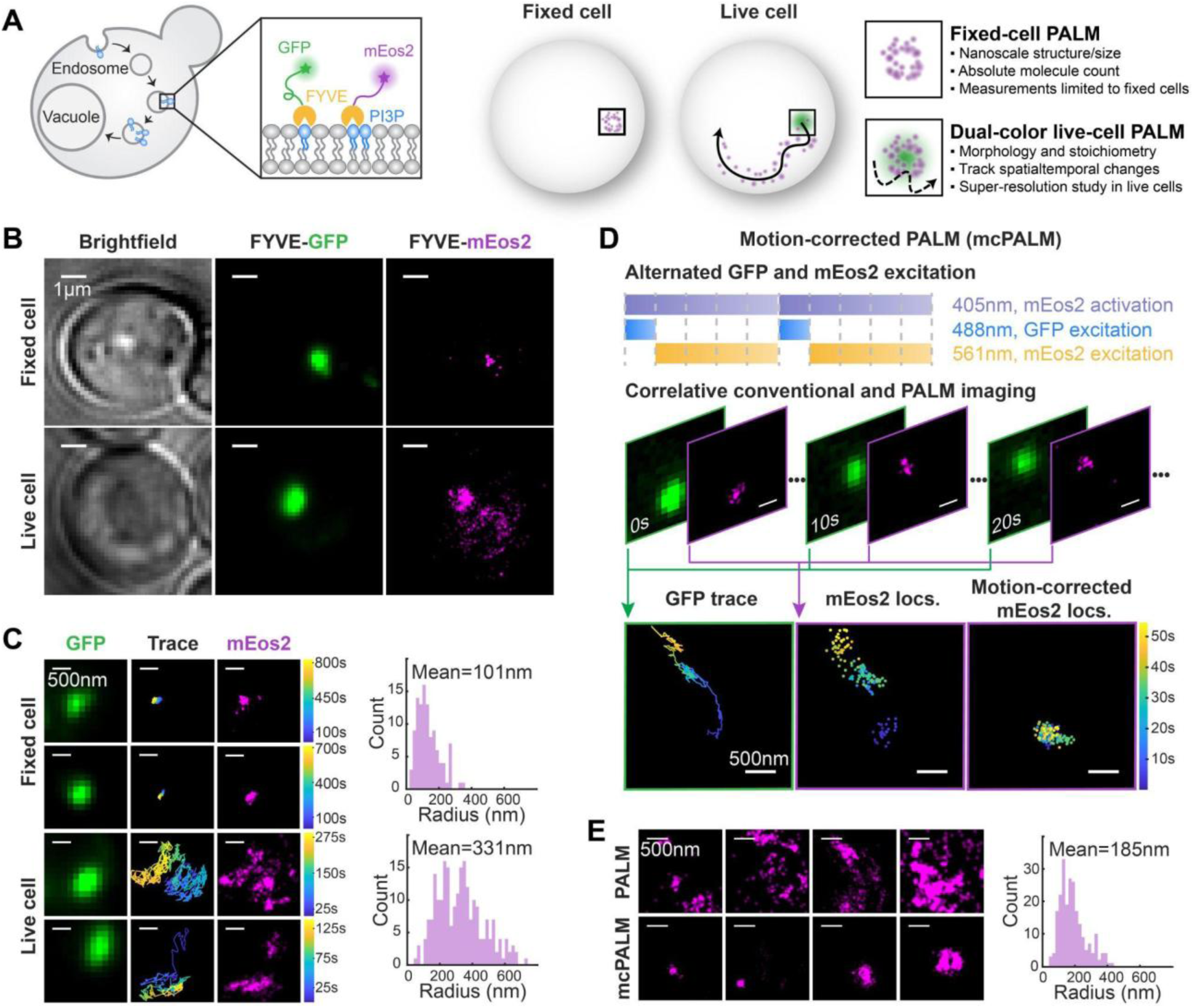
mcPALM corrects motion-induced spreading of single molecule localizations in living cells and improves endosome size measurement. **(A)** Left: schematic of the yeast endocytic pathway and fluorescent labeling strategy, in which the specific PI3P probe, a tandem FYVE domain, is tagged with both GFP and mEos2. Right: comparison between conventional PALM and dual-color PALM in fixed and live cells. Conventional PALM resolves nanoscale structure and molecule number but is limited to fixed cells, whereas dual-color PALM combines GFP-based tracking with mEos2 localization to enable super-resolution studies in live cells. **(B)** Top: in fixed cells, FYVE-mEos2 localizations in the PALM image colocalize well with FYVE-GFP endosomes in the conventional fluorescence channel. Bottom: in live cells, endosome motion smears FYVE-mEos2 localizations. Scale bar: 1 µm. **(C)** Examples of endosomes in the conventional fluorescence channel (left), their traces (middle) and raw single molecule localizations (right), which are smeared out in live cells (bottom). Scale bar: 500 nm. The effective endosome radius distribution in fixed cells has a mean of 101 nm calculated from 109 endosomes, whereas in live cells the spreading of mEos2 localizations along the endosome trajectories produces a broadened radius distribution with a mean of 331 nm calculated from 228 endosomes. **(D)** mcPALM workflow: a repeating shutter sequence contains continuous 405-nm photoactivation and alternating 488-nm GFP excitation for 1 frame and 561-nm mEos2 excitation for the next 4 frames. Motion correction is performed by subtracting the displacement of the GFP centroid from each mEos2 localization to obtain the corrected localization position. Scale bar: 500 nm. **(E)** After the mcPALM correction, FYVE-mEos2 localizations recover compact endosome structures. The motion-corrected effective radius distribution has a mean of 185 nm, a 44% decrease relative to the uncorrected measurement. Scale bar: 500 nm.

We first validated the dual-color probe in chemically fixed yeast cells, where endosome motion was eliminated. FYVE-mEos2 localizations colocalized with FYVE-GFP-positive endosomes, confirming that the conventional and PALM channels reported the same PI3P-positive compartments (Fig. 1B). Fixed-cell PALM revealed heterogeneous endosome morphologies ranging from smaller, donut-like structures to larger, elongated structures, similar to those in the previous fixed-cell study (Fig. 1B and 1C) (Puchner et al., 2013). To quantify endosome size, we calculated an effective radius from an anisotropic effective area enclosing 70% of mEos2 localizations relative to the localization center of mass (Supplementary Fig. S1 and Supplementary Materials). This analysis yielded a mean effective radius of 101 nm (Fig. 1C). This value is larger than the previously reported radius of 41 nm (Puchner et al., 2013), likely biased by our dual-channel workflow, which detected endosomes from the GFP channel, preferentially selecting brighter and thus larger endosomes, and also merged closely spaced late endosomal substructures or intraluminal vesicles into a single GFP-defined mask.

We next asked whether the same PALM-based structural measurements could be obtained in living cells. In live yeast cells, however, FYVE-mEos2 localizations were visibly smeared along the trajectories of moving endosomes over the long PALM data acquisition time, distorting reconstructed morphologies and inflating size measurements (Fig. 1B and 1C). The effective radius distribution broadened substantially, with an apparent mean of 331 nm (Fig. 1C). Because this broadening coincided with endosome displacement, we interpreted the increased radius as a motion-induced artifact rather than a reflection of the true physical dimension of the endosome. To correct this distortion, we developed motion-corrected PALM (mcPALM). This method leverages correlative dual-channel imaging with a repeating acquisition cycle containing five frames: one 488-nm GFP excitation frame to determine the endosome position in the conventional channel, followed by four 561-nm mEos2 excitation frames to collect localizations in the PALM channel, while weak 405-nm mEos2 photoactivation was applied throughout the cycle (Fig. 1D and Supplementary Video 1). During data processing, individual mEos2 localizations were then assigned to masks created by thresholding the GFP intensity of endosomes (Supplementary Fig. S1). For each mEos2 localization, we used the GFP-derived endosome trajectory as a motion reference and subtracted the corresponding frame-to-frame displacement from the mEos2 localization coordinates, thereby expressing the localizations in an endosome-centered coordinate system. This correction removed the apparent spreading caused by endosome motion and restored the compact endosome morphologies in live cells, similar to those observed in fixed cells (Fig. 1D and 1E). Quantitatively, mcPALM reduced the mean effective radius by 44%, from the uncorrected measurement of 331 nm to 185 nm, while narrowing down the radius distribution (Fig. 1E).

We also extensively quantified the additional localization uncertainty introduced by mcPALM. In conventional PALM, the uncertainty of each mEos2 localization is determined by photon number, finite pixel size, and background noise, and can be calculated using the Mortensen equation (Mortensen et al., 2010). In mcPALM, the localization uncertainty additionally includes contributions from localizing the GFP centroid, transforming localization coordinates between the spatially split GFP and mEos2 channels, and interpolating the GFP centroid position during the temporal offset between GFP and mEos2 excitation frames (Supplementary Materials and Supplementary Fig. S2). Combining these terms yielded a mean lateral localization uncertainty of 67 nm (Supplementary Materials and Supplementary Fig. S3), which represents reduced effective resolution compared with fixed-cell PALM, which has a mean lateral localization uncertainty of 29 nm (Supplementary Fig. S3), but remains well below the diffraction limit. These results establish mcPALM as a super-resolution microscopy technique in living cells that recovers nanoscale structural information from moving endocytic compartments, including compact localization distributions and size estimates, which otherwise cannot be reliably quantified by conventional PALM alone.

### Photoactivation-based correction improves single molecule counting of PI3P sites along the live-cell endosome maturation trajectory

In addition to resolving endosome size, mcPALM can quantify another key feature of endosome maturation: PI3P abundance. Since FYVE domains specifically bind to exposed PI3P sites, counting FYVE-mEos2 molecules provides an estimate of the accessible PI3P pool available for recruitment of effector proteins containing FYVE or PX domains.

Accurate molecule counting with mcPALM requires correction for two opposing sources of error. First, mEos2 blinking can lead to overcounting, as repeated localizations from the same molecule may appear as nonconsecutive events after transient transitions through short-lived dark states (Lee et al., 2012; Puchner et al., 2013). Because these events are expected to be proximate in space and time, we corrected this by merging mEos2 localizations within defined spatial and temporal thresholds, Δ*d* and Δ*t*, into a single molecule (Materials and Methods). Second, molecule numbers can be underestimated in live-cell imaging because endosomes or other moving fluorescent targets are trackable only for a limited duration. Endosomes can move out of the focal plane, and photobleaching in the reference channel can induce irreversible fluorescence loss, causing the recorded endosome trajectory to terminate before all mEos2 molecules have been detected during mcPALM acquisition (Fig. 2A).

**Figure 2.**
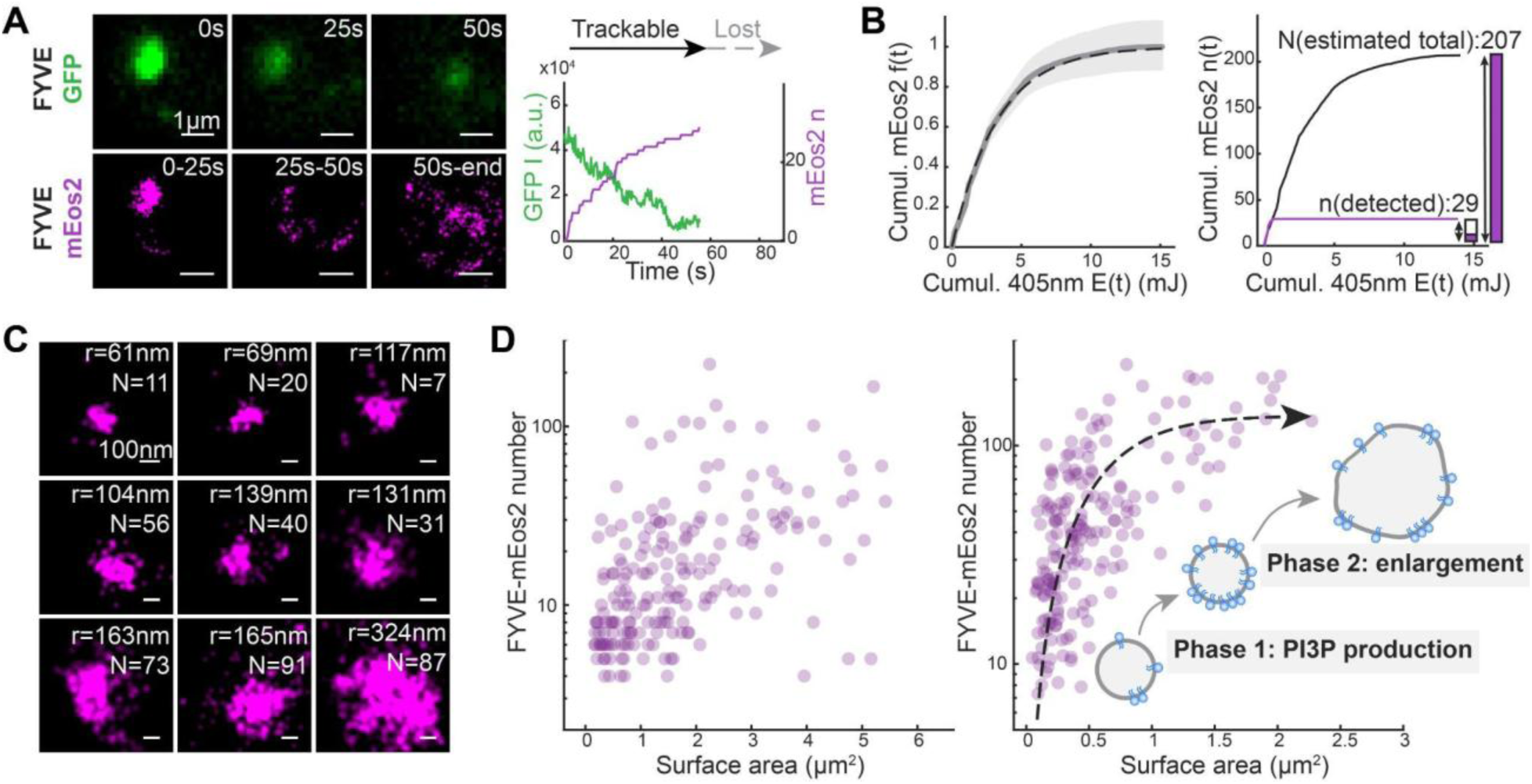
mcPALM-based measurements of endosome size and PI3P-bound FYVE-mEos2 numbers in live yeast cells reveal a two-phase maturation trajectory, consistent with results measured in fixed cells. **(A)** Short observation window and incomplete tracking limit mEos2 counting on moving endosomes. Top left: representative FYVE-GFP images (green) of a single endosome over time. Bottom left: corresponding motion-corrected FYVE-mEos2 localizations (magenta) accumulated in successive time windows. Right: progressively decreasing GFP intensity (green) due to bleaching and cumulatively increasing mEos2 number (magenta) from photoactivation. When GFP intensity falls to a low level, this endosome is lost and subsequent mEos2 localizations cannot be assigned correctly. Scale bar: 1 µm. **(B)** A stochastic photoactivation-based method to correct for undercounting of mEos2 molecule numbers from incomplete trajectories. Left: cumulative distribution function (CDF) of photoactivated mEos2 *f*(*t*), as a function of cumulative 405-nm laser energy *E*(*t*). The gray solid line represents the mean CDF from 11 movies, which is fitted to an exponential function shown as a dashed line. The shaded region represents the SEM across the 11 movies. Right: application of the calibration to the endosome in (A). Extrapolation based on the detected fraction allows estimation of the total mEos2 molecule count (N = 207) from the limited detection (n = 29). **(C)** Examples of endosomes from mcPALM spanning a range of motion-corrected sizes and corrected FYVE-mEos2 counts. Scale bar: 100 nm. **(D)** Left: relationship between apparent endosome surface areas and FYVE-mEos2 molecule numbers measured without motion correction and without molecule number correction shows no dependence. Right: motion-corrected endosome surface area plotted against calibrated FYVE-mEos2 numbers, which reports the number of accessible PI3P sites on individual endosomes. Data from 193 endosomes are shown. The dashed curve is an exponential guide to the eye. These data reveal a maturation sequence with two distinct phases: (1) PI3P production, characterized by a steep increase in PI3P content, followed by (2) size increase, at a saturated PI3P level.

To correct for this undercounting, we developed a correction method based on the stochastic nature of mEos2 photoactivation. In short, mEos2 photoactivation can be modeled as a Poisson process whose rate depends on the 405-nm laser power. Thus, the cumulative distribution function (CDF) of activated mEos2 molecules *f*(*t*) follows an exponential relationship with respect to the cumulative delivered 405-nm energy *E*(*t*) (Supplementary Materials and Supplementary Fig. S4). We therefore generated a *f*(*t*) - *E*(*t*) calibration curve, which is independent of the expression level of mEos2 (Fig. 2B and Supplementary Fig. S4). This calibration curve was then used to calculate the actual molecule number *N* for each tracked endosome by extrapolating the molecule number *n* detected between trace start time *t*_1_and disappearance time *t*_2_ to the point of 100% activation (Fig. 2B and Supplementary Materials). The statistical uncertainty of this corrected molecule number *N* was quantified using the standard deviation of the negative binomial distribution, providing an *n*- and *f*-dependent uncertainty estimate for the molecule count on each endosome (Supplementary Materials and Supplementary Fig. S4).

Thus, by simultaneously measuring motion-corrected sizes of endosomes and their photoactivation-calibrated FYVE-mEos2 molecule numbers, mcPALM allowed us to retrieve an endosome maturation trajectory in living cells, relating their surface area to the accessible PI3P content (Fig. 2C and 2D). This live-cell trajectory was consistent with both the previously reported fixed-cell PALM results and with our fixed-cell benchmarks analyzed using the same mcPALM pipeline (Supplementary Fig. S3). First, the number of accessible PI3P sites increased up to ∼150 on endosomes whose size remained relatively constrained, indicating an early phase dominated by PI3P production. In the second phase, endosomes continued to enlarge while the accessible PI3P number remained roughly constant, consistent with a transition from PI3P accumulation to membrane expansion.

We further evaluated the reliability of these measurements by color-coding the maturation trajectory according to the uncertainties in surface area and molecule number, and by comparing these patterns with fixed-cell analysis (Supplementary Materials and Supplementary Fig. S3–S4). In the trajectory color-coded by surface-area uncertainty and uncertainty-induced broadening bias, endosomes with similar PI3P levels but larger measured sizes tended to show higher size uncertainties and greater size inflation, suggesting that the measured uncertainty from error sources described in the last section at least partially contributes to the broader distribution compared to the fixed-cell benchmark, which indeed exhibited a narrower distribution (Supplementary Fig. S3). These results verify the robustness of mcPALM for extending quantitative PALM into living cells, enabling simultaneous quantification of structural sizes and absolute molecule numbers together with interpretable uncertainty estimates.

### mcPALM captures time-resolved transitions across mobility states of moving endosomes

In addition to nanoscale structure and molecule number, mcPALM also provides access to the real-time endosome trajectories which are not accessible from fixed-cell PALM. In living yeast cells, PI3P-marked endosomes displayed heterogeneous motion behaviors that could be grouped into different mobility classes, potentially suggesting the presence of distinct subpopulations with different transport functions. Only a small fraction of endosomes exhibited near-Brownian diffusion throughout the full observation window. Most endosome trajectories instead contained anomalous motion, including confined motion within various confinement sizes (Fig. 3A, traces 2–3), as well as intermittent directed transport (Fig. 3A, traces 4–5).

**Figure 3.**
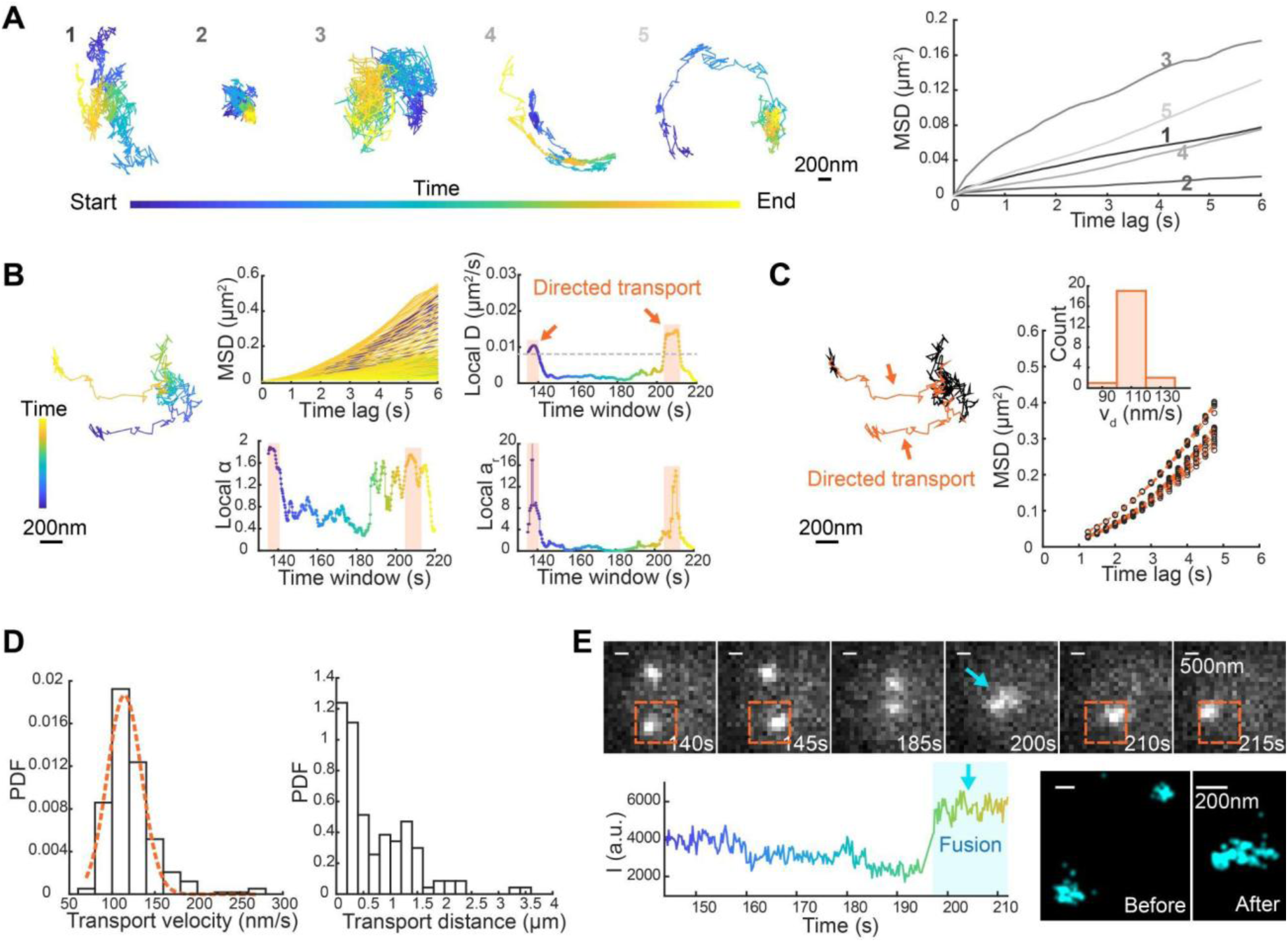
Directed transport and fusion dynamics of PI3P-positive endosomes revealed by local trace analysis. **(A)** Representative trajectories from five FYVE-positive endosomes. Trajectories are color-coded by time from start to end, and the corresponding MSD vs. time curves are shown on the right. Scale bar: 200 nm. **(B)** Sliding window analysis and identification of transient directed transport for one example endosome. Left: trace of the selected endosome, color-coded by time from start to end. Scale bar: 200 nm. The four panels on the right display features calculated within sliding windows along the trajectory: MSD, local diffusion coefficient D, local anomalous exponent ɑ and local directionality metric ar (angle ratio). **(C)** Analysis of directed transport segments in the same trajectory shown in (B). Left: segments classified as directed transport are highlighted in orange. Right: MSDs of directed transport segments fitted to the quadratic model. Only fits with *R*^2^ > 0.95 are shown. Inset: histogram of directed transport velocities *v*_*d*_ obtained from the quadratic fits. **(D)** Left: probability density function (PDF) of directed transport velocities *v*_*d*_ collected from 291 directed transport segments and identified across 193 endosomes, with *R*^2^ > 0.95. The distribution is fitted to a one-peak Gaussian distribution (dashed line) with a peak velocity of 114.44 nm/s. Right: distribution of transport distances for individual directed transport events. After merging temporally connected directed transport segments, they correspond to 117 directed transport events. **(E)** Time-lapse images of the same endosome analyzed in (B) and (C) showing an endosome fusion event. Top: FYVE-GFP images at the indicated time points. The orange dashed boxes mark the directed transport events in C, and the blue arrow indicates fusion. Bottom left: corresponding FYVE-GFP intensity of the tracked endosome over time, showing a sharp increase during fusion. Bottom right: FYVE-mEos2 mcPALM reconstructions before and after fusion. Scale bars are 500 nm in FYVE-GFP panels and 200 nm in super-resolution images.

Since individual endosomes often transitioned across different mobility states, the global mean squared displacement (MSD) fitting over the entire trajectory was insufficient to capture temporally varying dynamics. We therefore implemented a sliding window analysis to quantify local motion features, where we moved a window (T = 10 s) along each trajectory and performed MSD and step analyses on each segmented interval. From each segment, we extracted local parameters including the apparent diffusion coefficient *D*, the anomalous exponent *α*, and a directionality metric, *a*_*r*_, defined as the ratio of forward to backward steps (Izeddin et al., 2014). This approach enabled time-resolved visualization of mobility-state changes within individual endosome trajectories and allowed identification of transient directed transport (Fig. 3B).

Within each directed transport segment, we fitted the MSD to a quadratic directed transport model yielding the transport velocity *v*_*d*_, independently of random diffusive contributions (Fig. 3C). From all FYVE-positive endosomes with at least one directed transport component, the resulting transport velocities *v*_*d*_ had a peak of 114.44 nm/s, while transport distances peaked at a few hundred nanometers and extended into the micrometer range (Fig. 3D). This velocity is comparable in magnitude to many actin-dependent endosome velocities reported in previous studies using speed measurements and other endosomal markers (Toshima et al., 2006, 2016; Chang et al., 2005, 2003). Interestingly, many directed transport events followed nonlinear or curved paths, sometimes including sharp turns and bidirectional motion. Because many tracked endosomes moved along the vacuole, we could not determine whether this curvature reflected geometric confinement by the vacuole membrane or a property of the underlying transport mechanism. Nevertheless, these nonlinear paths are reminiscent of previously reported endosome movements driven by Las17p- and Arp2/3-dependent actin polymerization (Chang et al., 2005, 2003), which showed similar nonlinear trajectories and occasional sharp turns.

Additionally, we observed that some directed transport segments were associated with endosomal fusion or fission events (Fig. 3E and Supplementary Video 2). In these cases, live-cell mcPALM captured the dynamic transition in real time through the conventional channel while reconstructing super-resolved endosome morphologies before and after such events in the PALM channel. This highlights another advantage of mcPALM, that it can incorporate membrane remodeling processes and provide temporal super-resolution study on those dynamic events.

### An endosome motility shift with PI3P accumulation and size enlargement is revealed by mcPALM

Having shown that mcPALM can resolve transient mobility states within individual endosome trajectories, we next examined how endosome dynamics might change during their maturation, which is characterized by an increase in PI3P followed by an increase in size.

To obtain a trace-level descriptor of endosome motility, we averaged the local diffusivity *log*(*D*) and anomalous exponent *α* over all sliding-window segments within the same trajectory. To compare endosome motilities across maturation stages, we divided the PI3P-size maturation graph into three sections (Fig. 4A). Section 1 represents an early stage containing smaller endosomes with relatively low accessible PI3P. Section 2 contains endosomes with high PI3P content but relatively constrained size, while section 3 corresponds to large, mature endosomes with high, saturated PI3P content. The within-trace averaged diffusivity *log*(*D*) was significantly increased from Section 1 to sections 2 and 3, showing that PI3P accumulation is associated with increased endosome motility (Fig. 4A and 4B left). In contrast, the within-trace averaged anomalous exponent *α* did not differ significantly between sections 1 and 2, but decreased for large, mature endosomes (Fig. 4A and 4B right), suggesting that these enlarged endosomes are more likely to undergo confined motion although they retain high diffusivity.

**Figure 4.**
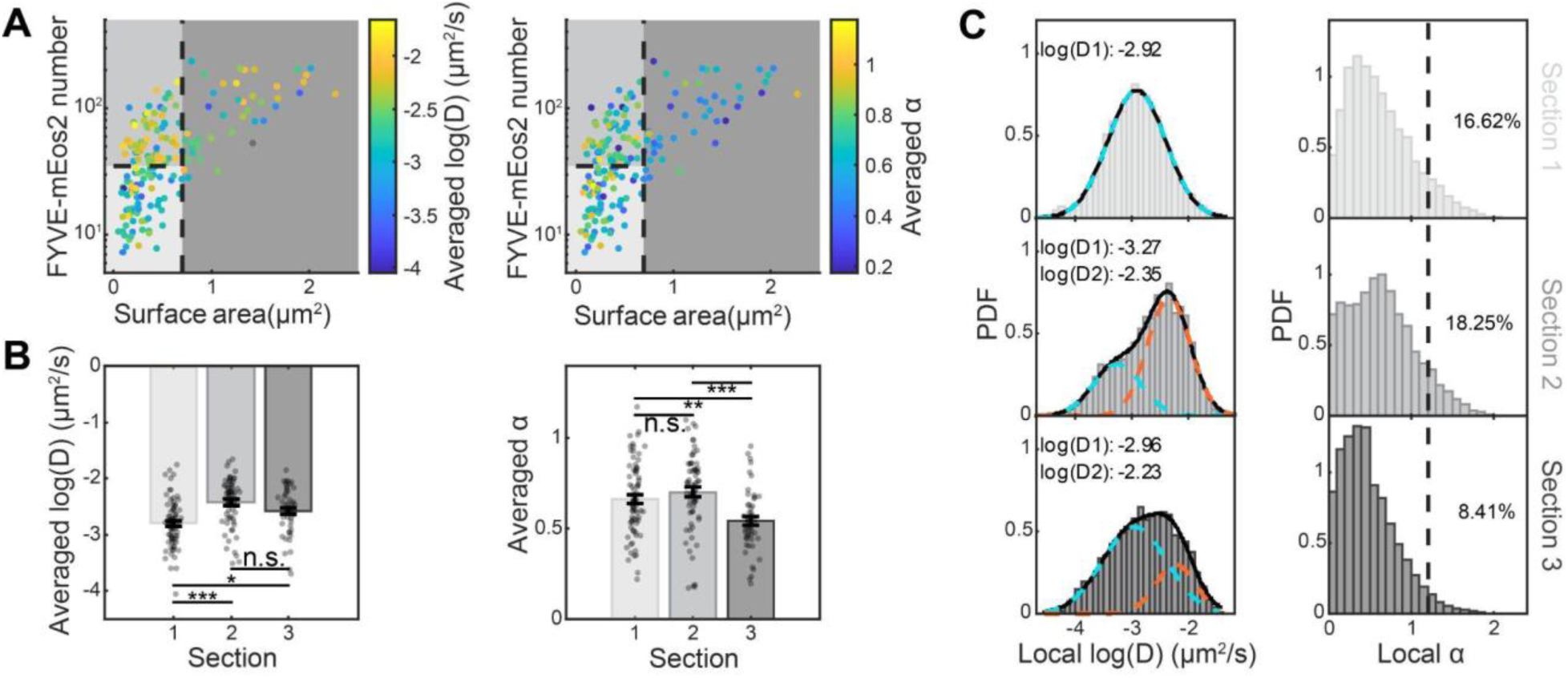
Bulk analysis of endosome dynamics reveals a motility shift across three stages of the PI3P-surface-area maturation trajectory. **(A)** The maturation landscape from Figure 2 color-coded by within-trace averaged log(D) (left) and anomalous exponent α (right). The dashed lines show the boundaries defining the three maturation sections (surface area = 0.7 µm^2^, FYVE-mEos2 number = 35). Colors are coded from light gray to dark gray for sections 1-3. **(B)** Bar plots of within-trace averaged log(D) (left) and anomalous exponent α (right) across three sections. Dots represent individual endosomes. Bars indicate mean ± SEM. Statistical significance is assessed by one-way ANOVA test plus Tukey’s Honestly Significant Difference; n.s. no significance; * p < 0.05; ** p < 0.01; *** p < 0.001. Cohen’s f values are 0.39 and 0.31. The numbers of endosomes in each section are 75, 67 and 48, respectively. **(C)** Probability density functions (PDFs) of local diffusivity log(D) (left) and local anomalous exponent α (right) from all trace segments within each section. In the log(D) distribution, fitting models with one, two or three peaks are selected by Bayesian information criterion (BIC) with penalty for model complexity. Section 1 is fitted by a single-peak Gaussian with log(D) = -2.92, whereas sections 2 and 3 are fitted by two-peak Gaussians, with log(D1) = -3.27 and log(D2) = -2.35 for Section 2, and log(D1) = -2.96 and log(D2) = -2.23 for section 3. In the α distribution, the fractions of segments with α > 1 are 16.62%, 18.25%, and 8.41% for Section 1, 2, and 3. The numbers of segments are 40921, 23247 and 23338 respectively.

The distributions of local *log*(*D*) and *α* values across all sliding windows within each section exhibited the same trend. Section 1 was dominated by a single slow diffusing population with local *D* centered at 10^-2.92^ µm^2^/s (Fig. 4C). In sections 2 and 3, the local *D* exhibited two-peak distributions, with a slower population below 10^-2.5^ µm^2^/s and a faster population above this threshold. The emergence of this fast diffusing population in PI3P-rich endosomes suggests that PI3P increase is accompanied by a more mobile state. However, although section 3 retained a mobile population, its anomalous exponent *α* distribution shifted toward lower values. Only 8.41% of segments in section 3 showed *α* > 1, compared with 16.62% and 18.25% in sections 1 and 2, respectively.

Together, these motility analyses revealed two coupled changes in endosome dynamics during maturation. First, PI3P accumulation is accompanied by increased diffusivity and may promote access to a more mobile state. The underlying reason for this increased motility remains unclear, but it may still be actin-associated (Chang et al., 2003, 2005) and involve recruitment of PI3P-dependent effector proteins that facilitate endosome transport. Second, later endosome enlargement is associated not simply with slower motion, but instead with increased confinement despite continued high diffusivity, potentially reflecting stronger interactions with or docking along the vacuole membrane. More broadly, this analysis demonstrates that live-cell mcPALM can simultaneously quantify nanoscale morphology, molecular composition, and motility changes, revealing how these features are linked in regulating organelle maturation.

### Dual-marker mcPALM maps PI3P-marked endosomes across Rab-defined maturation stages

A further advantage of mcPALM is that the conventional reference marker and the PALM probe do not need to be attached to the same molecule. Instead, they can label different proteins and report different molecular identities. As long as the two fluorophore signals partially or fully colocalize, motion correction can still be performed, and additional information can be extracted, such as the abundance of the second molecule and its relationship with structure size and dynamics. To demonstrate this capability and to relate PI3P-positive compartments to specific endocytic maturation stages, which are defined by the presence of specific Rab GTPases, we replaced the FYVE-GFP reference probe with (i) Vps21-GFP, the yeast Rab5 homolog (Horazdovsky et al., 1994; Gerrard et al., 2000) and (ii) Ypt7-GFP, the yeast Rab7 homolog (Cabrera et al., 2014; Wichmann et al., 1992). Vps21 marks early endosomes and regulates endosome-vesicle or endosome-endosome fusion through incorporating CORVET tethering complexes (Langemeyer et al., 2018; Peplowska et al., 2007; Markgraf et al., 2009), whereas Ypt7 associates with late endocytic compartments and promotes endosome-vacuole interactions by recruiting HOPS tethering complexes (Lachmann et al., 2011; Seals et al., 2000).

In the GFP channel, Vps21-positive endosomes were distributed throughout the yeast cell, and Ypt7 was enriched on the vacuole membrane but also appeared as bright puncta near the vacuole in some cells (Fig. 5A). For both markers, FYVE-mEos2 localizations showed substantial motion-induced spreading, but mcPALM correction recovered compact endosomal structures (Fig. 5B). Applying the same motion-correction and activation-calibration pipeline used for FYVE-GFP endosomes, we reconstructed a PI3P-size trajectory from Vps21- and Ypt7-labeled compartments (Fig. 5C). This characteristic trajectory was consistent with that obtained using FYVE-GFP (Fig. 2D) and with fixed-cell PALM measurements (Puchner et al., 2013). Along this trajectory, the Rab markers occupied distinct sections. Vps21-positive endosomes spanned a broad range of accessible PI3P levels from a few estimated FYVE-mEos2 molecules to ∼150 molecules, while remaining relatively constrained in size. In contrast, Ypt7-positive endosomes were associated with high PI3P content and displayed a broader size range extending to larger structures. These observations are consistent with the known associations of these Rab GTPases with different endosome maturation stages and validate mcPALM for use with two different markers.

**Figure 5.**
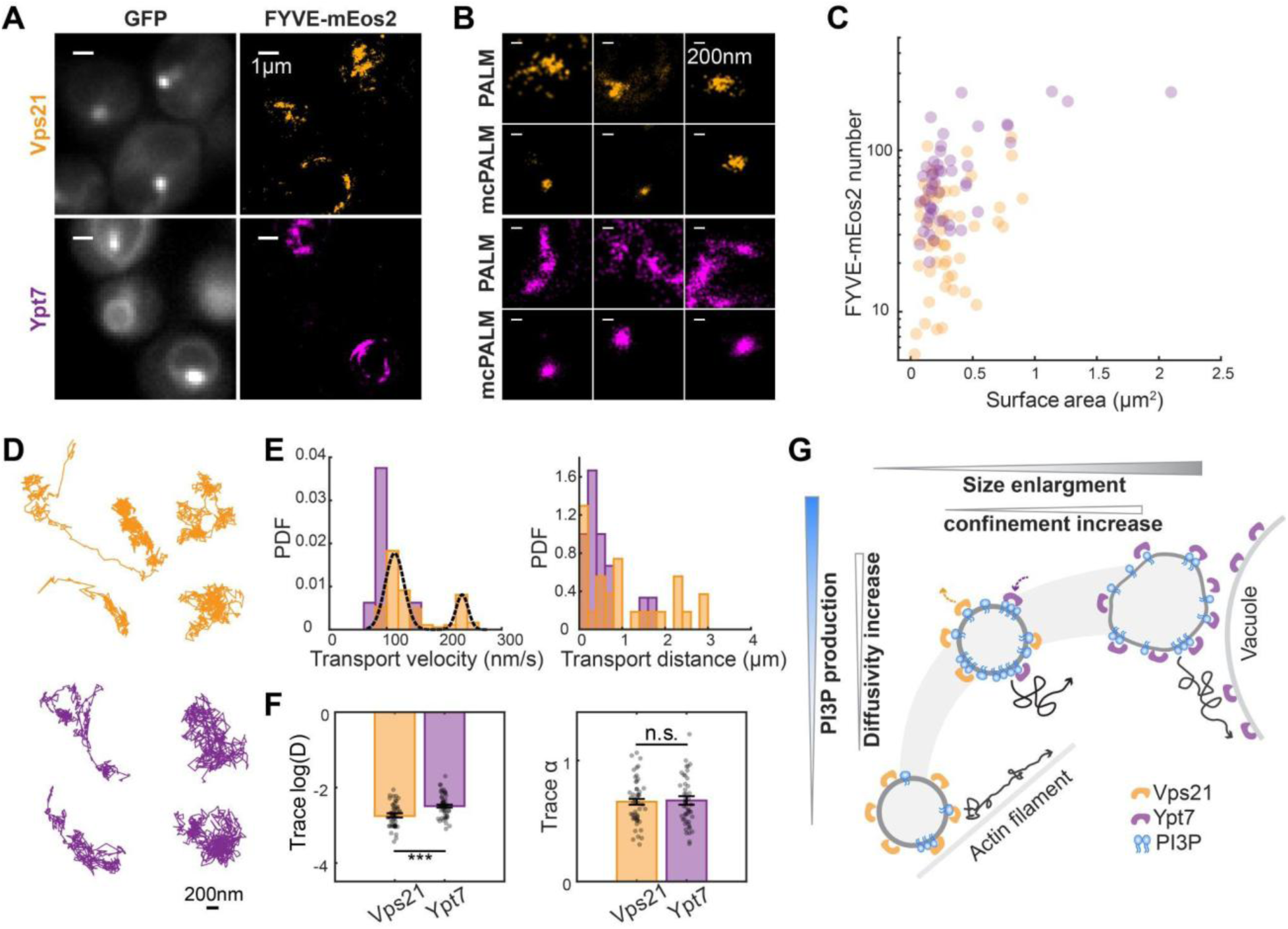
Vps21-positive (orange) and Ypt7-positive (magenta) endosomes show distinct structural and dynamic properties following the maturation trajectory. **(A)** Fluorescence images of yeast cells expressing Vps21-GFP and Ypt7-GFP together with FYVE-mEos2 localizations. Scale bar: 1 μm. **(B)** Examples of PALM and mcPALM images of Vps21-positive and Ypt7-positive endosomes. Scale bar: 200 nm. **(C)** Relationship between motion-corrected endosome surface area and accessible PI3P content, represented by corrected FYVE-mEos2 molecule numbers. This relationship is consistent with the maturation trajectory in Fig. 2. Ypt7-positive endosomes generally contain higher FYVE-mEos2 numbers and extend to larger sizes compared to Vps21-positive endosomes. Data are from 50 Vps21-positive endosomes and 50 Ypt7-positive endosomes. **(D)** Representative trajectories of Vps21-positive and Ypt7-positive endosomes. Scale bar: 200 nm. **(E)** Probability density functions (PDFs) of directed transport properties. Left: distributions of directed transport velocities calculated from the quadratic fits to segmental MSDs, as described in Fig. 3. 93 segments from 50 Vps21-positive endosomes and 8 segments from 50 Ypt7-positive endosomes are included in the statistics after retaining fits with *R*^2^ > 0.95. The dashed curve over the Vps21 directed velocity distribution shows a two-peak Gaussian fit, with peaks at 113.56 nm/s and 231.10 nm/s. Right: PDFs of directed transport distances for individual directed transport events. All segments from Vps21-positive endosomes are merged to 27 directed transport events, and segments from Ypt7-positive endosomes are merged to 15 directed transport events. **(F)** Quantification of trajectory dynamics reveals a motility shift across different maturation stages. Shown are within-trace averaged diffusivity log(D) (left) and anomalous exponent α (right). Statistical significance is assessed by one-way ANOVA test, with n.s. no significance, * p < 0.05, ** p < 0.01, *** p < 0.001. Cohen’s d values are -0.82 and -0.04, respectively. As in (C), 50 Vps21-positive endosomes and 50 Ypt7-positive endosomes are included in the statistics. **(G)** Schematic of the dynamics-integrated endosome maturation landscape. As PI3P accumulates, endosome diffusivity increases. Then Vps21-to-Ypt7 conversion happens at high PI3P level. At later stages, endosomes enlarge and become more confined.

Dynamic analysis further distinguished the two populations. Vps21-positive endosomes exhibited diverse trajectories, including diffusive, confined, curved, and occasionally linear directed transport paths (Fig. 5D and Supplementary Video 3). Ypt7-positive compartments also transitioned among different motion behaviors, but their trajectories appeared mainly associated with the vacuolar surface (Fig. 5D and Supplementary Video 3). Analysis of directed transport segments showed that Vps21-positive endosomes exhibited a two-peak distribution of directed transport velocities *v*_*d*_, with one peak at 113.56 nm/s, similar to FYVE-GFP-labeled endosomes, and a distinct, faster peak at 231.10 nm/s, derived primarily from straighter, more linear transport paths (Fig. 5E). In contrast, Ypt7-positive structures showed a narrower transport velocity distribution centered near 100 nm/s and generally shorter transport distances. These results suggest that Vps21-labeled early endosomes are more likely to undergo faster and more linear directed transport, potentially driven by a distinct actin-dependent transport mechanism that can support such longer-range transport. This is in line with previous reports that early endosomes depend on formin-dependent actin polymerization and can attach to actin clumps that are coupled to actin cable flow (Toshima et al., 2016; Huckaba et al., 2004; Girao et al., 2008).

Interestingly, despite their more prominent directed transport behavior, Vps21-positive endosomes had a lower within-trace averaged diffusivity *log*(*D*) than Ypt7-positive structures (Fig. 5F). This is consistent with the motility shift along the maturation trajectory observed using FYVE markers, where higher PI3P levels were associated with increased diffusivity (Fig. 4). Local diffusivity and step distributions also support this difference. Most Vps21-positive trajectory segments occupied a slower diffusivity region centered at 10^-2.83^ µm^2^/s, whereas a faster diffusing phase peaking at 10^-2.27^ µm^2^/s was more prominent among Ypt7-positive segments (Supplementary Fig. S5A). Moreover, step sizes in T = 1 s intervals from Ypt7-positive endosomes showed a larger fraction of >150 nm than those from Vps21-positive ones (Supplementary Fig. S5B). However, Vps21-positive endosomes showed greater within-trace heterogeneity in both diffusivity and anomalous exponent, as measured by the standard deviation of local values within each trace (Supplementary Fig. S5C), showing agreement with the observation that early endosomes more frequently switch between slower, fusion- or fission-associated motion and faster, actin-associated directed transport (Supplementary Video). Thus, the lower average diffusivity of Vps21-positive endosomes may reflect a greater fraction of time spent in slow-moving, interaction-heavy periods. By contrast, Ypt7-positive structures showed higher average diffusivity but lower heterogeneity, meaning more persistent motility.

The within-trace averaged anomalous exponent *α* did not differ significantly between Vps21- and Ypt7-positive populations (Fig. 5F), and the fraction of segments with *α* > 1 decreased only slightly from 16.21% in the Vps21 population to 13.30% in the Ypt7 population (Supplementary Fig. S5A). However, the directionality metric *log*_2_*a*_*r*_ became more negative for larger step sizes of Ypt7-positive structures (Supplementary Fig. S5B), suggesting the presence of long-range confinement, likely reflecting tethering and docking along the vacuole membrane.

Together, these results show that dual-marker mcPALM can be extended beyond same-molecule labeling to different PALM and conventional fluorescence marker proteins. This approach positions PI3P-positive endocytic compartments within a Rab-defined, dynamics-integrated maturation landscape (Fig. 5G), where Vps21-positive endosomes occupy the PI3P-increasing stage with stronger directed transport and greater within-trace variability, whereas Ypt7-positive compartments occupy the PI3P-rich, enlarged stage with higher and more persistent diffusivity but stronger confinement near the vacuole. Thus, this marker-flexible strategy allows us to link second-marker identity and abundance to nanoscale structure, first-marker stoichiometry, and dynamic behavior in living cells.

## Discussion

Quantitative PALM has often been restricted to chemically fixed cells or immobile structures to prevent motion-induced image distortion and to preserve high spatial resolution. In this work, we developed mcPALM as a super-resolution framework that bridges quantitative PALM with live-cell imaging. By pairing PALM with a conventional fluorescence reference channel, mcPALM tracks target motion during data acquisition, corrects motion-induced spreading of single-molecule localizations, and reconstructs nanoscale morphology while preserving real-time dynamic information. Although live-cell mcPALM has higher empirical localization uncertainty than fixed-cell PALM, its effective resolution remains below the diffraction limit and still supports biologically informative single-molecule measurements. Thus, mcPALM is useful for quantitatively analyzing moving intracellular structures whose biological functions depend on both molecular organization and dynamic behavior, which therefore need to be measured together.

We tested the robustness of quantitative mcPALM by benchmarking it against the well-characterized endosome maturation pathway in yeast. mcPALM recovered compact morphologies of moving endosomes and substantially reduced their mean effective radius by 44%, validating its ability to correct motion-induced distortion. However, the corrected live-cell radius remained larger than that measured in fixed cells, reflecting several limitations of the current implementation. First, live-cell mcPALM accumulates additional localization uncertainties from the conventional reference channel. We therefore provide guidance on uncertainty quantification from multiple error sources and on potential optimization for user-defined imaging conditions. For instance, if this framework is applied to faster targets, reducing the time offset between the conventional-reference and PALM acquisition frames, or using simultaneous acquisition, would improve localization precision by decreasing the interpolation error. On the other hand, reducing this uncertainty comes with a trade-off. More frequent reference-channel excitation increases photobleaching, thereby consuming the finite photon budget more quickly and shortening the available tracking duration. Thus, the excitation sampling rate should be carefully optimized by balancing target motion against reference-channel photostability. More photostable fluorophores, such as the newly developed StayGold (Hirano et al., 2022), may further improve tracking duration and precision. Second, the current requirement to detect endosomes in the conventional channel biases the analyzed population toward larger endosomes with sufficient GFP signals, and rotational motion or shape change that mcPALM currently cannot account for also increases the time-averaged radius of non-spherical structures. Despite these limitations, many measured endosome sizes were still below the optical diffraction limit, and the relative comparison between different endosome populations remained informative as they are subject to similar size-increasing artifacts. Nevertheless, future improvements could consider incorporating correction for shape change and rotational motion, as well as using less biased reference markers.

While we demonstrated mcPALM using PI3P-regulated endosome maturation in yeast, this method can be extended to many other molecular markers and biological systems. The conventional reference fluorophore and the PALM-compatible fluorophore do not need to label the same molecule, as long as their signals colocalize sufficiently to define the same moving structure. This allows mcPALM to quantify relationships between different molecular components, as demonstrated here by mapping Rab GTPases to distinct PI3P-marked endosomal populations. We have also previously applied a similar motion-correction strategy to telomere clusters in live tumor cells, where it enabled reliable separation of bound and unbound dCas9/MCP complexes, supporting the broader applicability of mcPALM (Mehra et al., 2022). With appropriate optimization of imaging conditions, potential future applications include vesicles, organelle contact sites, membrane-associated signaling clusters, protein assemblies, and other mobile subcellular structures. Beyond that, incorporating axial tracking and three-dimensional localization could further improve precision and extend mcPALM to volumetric live-cell super-resolution imaging.

Single molecule counting is another unique capability of PALM, providing quantitative information beyond morphology. Here we developed a photoactivation-based correction to improve single molecule counting in live cells. This correction addresses molecule undercounting caused by incomplete trajectories of moving targets. Since the fraction of photoactivated molecules follows a reproducible saturating exponential rise with respect to the cumulative photoactivation energy, molecules detected during a finite observation window can be treated as a known fraction of the total molecule pool and extrapolated to estimate the actual molecule number. This strategy is applicable to many live-cell scenarios in which the target remains observable for only a limited time, such as when targets leave the field of view, move out of focus, or lose the reference-channel signal due to marker turnover or photobleaching.

Several factors remain important for reliable counting. First, proper fluorophore choice is critical. Single molecule counting is best suited for irreversibly photoactivatable or photoconvertible fluorophores that eventually enter a photobleached state, such as mEos2 used here and Dendra2, reducing complications from repeated photoactivations. Second, the photoactivation-based correction assumes steady state, meaning the molecule number on each tracked structure remains approximately constant during the observation window. Third, short traces or low-detection events produce larger uncertainty for this live-cell correction. We therefore provide guidelines to evaluate statistical uncertainty and apply filtering to retain traces with sufficient detected molecules while excluding poorly constrained estimates. Additional unaddressed considerations include spatially varying photoactivation/photophysics and limited detection efficiency. For example, laser illumination is not strictly uniform across the field of view, and ∼40% of mEos2 molecules are undetectable under PALM imaging conditions (Puchner et al., 2013) due to reasons including incomplete photoconversion and incomplete protein maturation. Future improvements could focus on performing position-dependent correction and on accounting for the non-detectable fraction.

By combining mcPALM with photoactivation-based counting correction, we simultaneously quantified three critical parameters of endosome maturation: endosome size mostly at or below the diffraction limit, molecule content of distinct lipids or proteins defining maturation stages, and diverse endosome dynamics. These measurements allowed mcPALM to successfully recover the known endosome maturation trajectory observed by fixed-cell PALM, characterized by an initial phase of PI3P production followed by an increase in endosome size. Importantly, live-cell mcPALM further allowed this structural and molecular trajectory to be integrated with dynamic information of moving endosomes, building a dynamics-integrated maturation landscape (Fig. 5G). From this landscape, we showed how mcPALM links time-resolved motion analysis with structural and stoichiometric measurements. Two aspects of heterogeneous endosome dynamics were extracted: transient directed transport and average motility. First, directed transport showed stage-dependent trends. Early-stage Vps21-positive endosomes exhibited a distinct fast, linear transport mode (*v*_*d*_ > 200 nm/s), for which formin-dependent actin polymerization is a strong candidate mechanism, as also in previous actin inhibitor studies (Toshima et al., 2016). Slower, curved directed transport (*v*_*d*_ ≈ 100 nm/s) was more broadly observed in other endosome stages and may involve Las17p- and Arp2/3-dependent actin polymerization, as reported for Ste2-labeled endosomes with similar trajectory properties (Chang et al., 2005, 2003; Girao et al., 2008). However, assigning correct force-generating mechanisms to each transport class will require direct perturbation of candidate actin pathways, for example using specific actin polymerization inhibitors, in future work. Second, average motility then showed a complementary maturation-dependent trend. PI3P accumulation was associated with the emergence of a faster diffusive state, whereas later endosome enlargement was accompanied by stronger spatial confinement near the vacuole. Consistent with this, although early-stage Vps21-positive endosomes exhibited faster transient directed transport, their average diffusivity was lower, likely because they spent a large fraction of time on slower mobility states during fusion and fission events. In contrast, PI3P-rich Ypt7-positive compartments displayed higher average diffusivity with less variability, in agreement with their more persistent motion along the vacuole. Together, these results indicate that endosome maturation involves not only changes in size, PI3P content, and Rab identity, but also coordinated shifts in dynamic behavior.

Beyond measuring motion, mcPALM also provides access to other temporally varying structural and molecular events that are not observable in fixed cells, such as membrane remodeling, organelle growth, and changes in molecular composition. We leveraged this capability and resolved endosome structures before and after fusion by dividing endosome trajectories into shorter temporal windows for analysis. While shorter windows better capture such dynamic changes at higher temporal resolution, they also lower localization sampling density and thereby reduce the effective spatial resolution according to the Nyquist criterion. Thus, the optimal window size should be chosen based on the timescale of the biological event and the number of single-molecule localizations required for sufficient spatial resolution, for example as estimated by Fourier ring correlation (Nieuwenhuizen et al., 2013; Banterle et al., 2013).

The findings in the endosome maturation system highlight the central strength of mcPALM: it links nanoscale morphology, molecular composition, and dynamics and reveals how these features are coupled in regulating biological processes and functions. In summary, mcPALM along with the photoactivation-based correction provides a practical framework for expanding the power of quantitative super-resolution imaging to moving objects in live cells. The pipeline combines automated motion correction, size quantification, molecule counting, dynamics analysis, and quantitative uncertainty estimation, thereby gaining rich and biologically meaningful structural, molecular, and dynamic information.

## Materials and Methods

### Yeast preparation

All yeast strains used in this study were derived from the W303 background and have been described previously (Puchner et al., 2013). Yeast strains were maintained on YPD plates at 4 °C. For imaging experiments, yeast cells were incubated in 2 mL synthetic complete dextrose (SCD) medium and grown overnight at 30 °C with shaking at 270 rpm. The next morning, cell cultures were diluted into fresh SCD medium to an optical density (OD) of ∼0.2 and grown for ∼4 hr under the same conditions until they reached the exponential phase, with an OD of 0.5–1.0. Eight-well chambered borosilicate coverglasses (C8-1.5H-N; Cellvis) were prepared by incubating each well with 100 µL of 0.8 mg/mL sterile Concanavalin A (ConA) for 0.5–1 hr followed by three washes with deionized water (DI water; Millipore). For live-cell imaging, 30 µL of yeast cell culture was mixed with 270 µL fresh SCD medium in each well and allowed to settle on the coated coverglass for 20–30 min prior to imaging. For fixed-cell imaging, 2 mL of cell culture grown to the exponential phase was harvested by centrifugation for 2 min at 2500 rpm (Mund et al., 2014). Cells were then resuspended in 100 µL DI water and added into the chamber to fully cover the glass surface. After 15–20 min of settling, DI water was completely removed and replaced with 100 µL of 4% formaldehyde solution. Cells were fixed for 15 min at room temperature with gentle shaking at 135 rpm. The formaldehyde solution was then replaced with 100 µL phosphate-buffered saline (PBS), and this wash was repeated twice. After the final wash, the PBS volume was adjusted to 300 µL to prevent evaporation during imaging and to maintain consistency with live-cell imaging conditions.

### Microscope setup

Fluorescence imaging was performed on a Nikon Ti-E inverted microscope (Eclipse Ti-E) equipped with a Perfect Focus System. Three continuous-wave lasers at 405, 488, and 561 nm were used for mEos2 photoactivation, GFP excitation, and mEos2 excitation, respectively (OBIS-CW; Coherent). The laser beams were aligned, expanded, and focused onto the back focal plane of a Nikon CFI Apo 100x Oil-immersion objective (N.A. 1.49) to illuminate the field of view. Laser illumination was controlled using a programmed shutter sequence that periodically turned lasers on and off. For correlative PALM and conventional fluorescence imaging, we used a five-frame acquisition cycle consisting of one 488-nm frame for GFP excitation and conventional imaging, followed by four 561-nm frames for mEos2 excitation and PALM imaging. The 488-nm laser power was 0.07–0.7 W/cm², and the 561-nm laser power was 0.5–1 kW/cm². During acquisition, the 405-nm laser was kept on for mEos2 photoactivation and was gradually increased from 0.1 to 2 W/cm².

Fluorescence emission was separated from the excitation illumination background using a quad-band dichroic mirror (zt405/488/561/640rdc; Chroma). The emission was then split into two channels by a long-pass dichroic beamsplitter (T562lpxr BS; Chroma), generating a green conventional fluorescence channel and a red PALM channel. Each channel was further filtered by band-pass filters (ET525/50 and ET595/50; Chroma). Images were acquired at 20 Hz using an EMCCD camera (iXon Ultra 897; Andor), which was cooled to −70 °C with an EM gain of 30. Movies were recorded using HAL4000 software (Zhuang lab, Harvard) and saved as DAX files.

### Motion correction pipeline

#### 1. Single molecule detection in the PALM channel (the mEos2 channel)

mEos2 single molecule detection and localization in the PALM channel were performed using Insight3 (Zhuang lab, Harvard). In each PALM frame, candidate mEos2 fluorescence spots were fitted with a 2D Gaussian function in a 7 × 7 pixel region of interest (ROI) at a pixel size of 160 nm. Fitted localizations were retained if they had a peak height ≥ 74 photons and a width between 200 and 700 nm. All localization information, including x and y coordinates, frame number, width, intensity, and other relevant parameters, was saved as a binary file for further analysis in MATLAB.

#### 2. Endosome detection in the conventional channel (the GFP channel)

Endosome detection and downstream motion correction were automated in MATLAB. GFP frames were first denoised using singular value decomposition (SVD) with a sliding time window of 5 frames and a sensitivity of 0.1. The denoised GFP frames were used only to improve the signal-to-noise ratio and increase the sensitivity of endosome detection. Each denoised frame was then thresholded using the built-in MATLAB function adaptthresh based on local GFP intensity to generate binary masks and boundaries for individual endosomes. The sensitivity parameter was adjusted between 0.16 and 0.26 for different GFP markers and kept constant for each marker, while the local neighborhood size was fixed at 7 pixels. Only masks with an area ≥ 6 pixels were included in the downstream analysis to exclude isolated noisy pixels and small background fluctuations.

Endosome centroids were determined using two approaches depending on mask size, summed GFP intensity and whether the GFP intensity profile could be reliably fit as a Gaussian PSF. For smaller endosomes with a mask area < 49 pixels, corresponding to the 7 × 7 pixel ROI used for single-emitter PSF detection, centroids were obtained by 2D Gaussian fitting in Insight3. For larger endosomes with a mask area ≥ 49 pixels, as well as endosomes that were not identified as valid Gaussian PSFs in Insight3 due to either low intensity or the GFP intensity profile deviating from the Gaussian model, centroids were calculated as the intensity-weighted center of mass of all pixels within the thresholded boundary. This calculation was performed using the undenoised, background-subtracted GFP image to avoid potential bias introduced by SVD denoising. Additional details of the GFP centroid determination procedure and its uncertainty analysis are listed in the Supplementary Materials.

Before motion correction, endosome traces were generated using a nearest-neighbor linking algorithm. Endosomes detected in different frames were linked if the temporal gap between two detections was less than 5 s and the spatial separation between them was less than 400 nm. This allowed endosomes to be linked across short dark periods, which can occur when GFP signals are temporarily weak, noisy or out of focus during threshold-based detection. The linking distance cutoff was then recalculated as the 99th-percentile of all step lengths from the initially generated traces, and tracking was repeated using this optimized cutoff. Only traces longer than 20 steps were retained for further analysis to exclude short traces with unreliable motion parameter fits. Next, because GFP excitation frames were acquired intermittently in the shutter sequence, endosome centroid positions were linearly interpolated between adjacent GFP excitation frames. Endosome boundaries were similarly interpolated by shifting the boundary from the previous GFP frame according to the interpolated centroid displacement. The rationale for the linear interpolation and the associated interpolation uncertainty analysis are provided in the Supplementary Materials.

#### 3. Channel transformation

Since the mEos2 and GFP emission channels were spatially split, a set of transformation equations was generated for each day of imaging to precisely map mEos2 localization coordinates in the PALM channel to (x′, y′) in the conventional channel. To determine the transformation equations, as described previously (Mancebo et al., 2020), we used a coverslip loaded with 20 µL of 130 µM fluorescein (Sigma-Aldrich) and imaged an 8 x 8 dot array projected onto the coverslip by a digital mirror device (DMD) under 405-nm illumination. Since each dot had a size below the diffraction limit, its fluorescence intensity profile was treated as a single-emitter PSF and localized via Insight3 in both GFP and mEos2 channels. The coordinates of each dot in both channels were then used to determine coefficients of the third-order polynomial transformation equations. The empirical uncertainty analysis for the channel transformation process is provided in the Supplementary Materials.

#### 4. Motion correction and size measurement

The transformed mEos2 localizations located within an endosome boundary in the corresponding frame were assigned to that endosome. Only endosomes with more than two mEos2 localizations were saved for downstream analysis to ensure reliable size and molecule number measurements. Each assigned mEos2 localization was then motion-corrected by subtracting the centroid displacement of the corresponding endosome measured in the GFP reference channel, thereby expressing the localization in an endosome-centered coordinate system. The resulting localization uncertainty after motion correction is discussed in the Supplementary Materials.

The effective radius of each endosome was calculated using an anisotropic effective area that enclosed 70% of mEos2 localizations around the center of mass, to reduce the influence of the false-positive localizations near the GFP-defined mask boundary. The corresponding surface area was estimated from the effective radius by assuming spherical geometry. Further details of the size measurement and its error analysis are described in the Supplementary Materials.

#### 5. Single molecule counting

As discussed in the Results section, live-cell molecule counting required blink correction and photoactivation-based correction. Blink correction for FYVE-mEos2 localizations was performed by grouping repeated localizations using two parameters: a maximum linking-distance threshold Δ*d* and a maximum dark-time threshold Δ*t*. Both thresholds were chosen based on the short-distance and short-time correlation peak associated with blinking events from the same molecule in the spatio-temporal cross-correlation analysis, as detailed in the Supplementary Materials. For fixed cells, the linking-distance and dark-time thresholds were set to 80 nm and 3 s, similar to values previously used for mEos3.2 (Banerjee et al., 2023). For live cells, the linking-distance and dark-time thresholds were set to 180 nm and 3 s. A custom MATLAB function was then used to compute the photon-weighted average position from the linked localizations.

The blink-corrected molecule numbers were further corrected using a photoactivation-based correction to account for undercounting caused by the finite observation time of endosomes during live-cell imaging (Figure 2 and Supplementary Materials). This correction was based on the cumulative photoactivation laser energy. A calibration curve was generated by averaging curves of cumulative detected mEos2 fraction vs. delivered photoactivation energy from all movies acquired on the same day. This calibration curve was then applied to extrapolate the detected fraction of mEos2 molecules on each endosome to 100% for estimating the underlying molecule numbers. All analyses here were performed using custom MATLAB functions. More details about this photoactivation-based molecule-counting correction are also provided in the Supplementary Materials.

### Endosome diffusion analysis

#### 1. MSD and step analysis

For the endosome traces obtained as described above, the mean squared displacement (MSD) was calculated by averaging the squared displacements over different time intervals. The diffusion coefficient *D* of each endosome trajectory was then calculated by fitting the MSD to a linear line according to the two-dimensional Brownian diffusion MSD model with a localization error term:

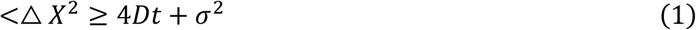

where *D* is the diffusion coefficient and *σ* is the localization error. Since many endosomes exhibited more complex motion than simple Brownian diffusion, the MSD was also fitted to a power-law model to determine the value of anomalous exponent *α*:

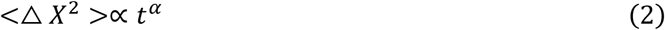

where *α* characterizes the deviation from random diffusion. An *α* value close to 1 indicates Brownian motion, whereas *α* > 1 suggests superdiffusive or directed motion and *α* < 1 suggests subdiffusive or confined motion.

In parallel with MSD analysis, we also performed step analysis as an additional measure of endosome motion. The average speed *v* was computed as the net distance *Δd* between the start and last point divided by the elapsed time *Δt*:

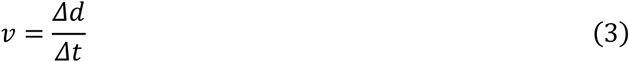

We further defined a step angle ratio *a*_*r*_ as a directionality metric inspired by (Izeddin et al., 2014):

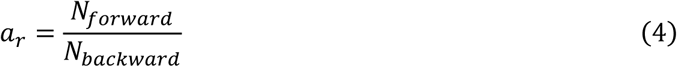

where *N*_*forward*_ and *N*_*backward*_ denote the numbers of forward and backward steps, respectively. Turning angles were defined between consecutive displacement vectors and constrained to the range of [0°, 180°]. Forward steps were defined as consecutive steps over a T = 1 s interval with turning angles < 40° and backward steps were defined as those with turning angles > 140°. A larger *a*_*r*_ indicates stronger directed transport, while a smaller *a*_*r*_ indicates more prominent confinement. We also calculated *log*_2_*a*_*r*_ for easier interpretation, so that a value of zero means balanced or random orientation, whereas positive values indicate directed transport-dominated motion and negative values indicate confinement-dominated motion.

#### 2. Sliding window analysis

To quantify local changes in endosome dynamics, each trajectory was divided into shorter intervals using a T = 10 s sliding window. MSD and step analyses were then performed within each window to extract local dynamic parameters. For directed transport analysis, individual segments were classified as directed transport if they had a local diffusion coefficient *D* ≥ 0.008 µm^2^/s and either a local anomalous exponent *α* ≥ 1.5 or a local step angle ratio *a*_*r*_ ≥ 10, reflecting the expectation that directed transport segments exhibit increased motility and stronger forward-step directionality. The MSD of each segment identified as directed transport was fitted to the quadratic model to extract the directed transport velocity *v*_*d*_:

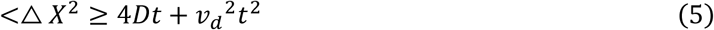

and only fits with *R*^2^ > 0.95 were included in the statistical analysis of *v*_*d*_ to minimize the influence of poorly fitted directed transport segments. For bulk motility analysis, all local diffusivity log (*D*) from all segments within a single trace were averaged to obtain the overall motility of that endosome. The same procedure was applied to the anomalous exponent *α*. The standard deviation of these values within each trace was used to represent the motility heterogeneity of that endosome.

## Supplementary Materials

Additional experimental details, analytical methods and error analyses are listed in the Supplementary Materials. Supplementary videos are also included.

## Data Availability

All raw microscopy movie files and yeast strains used in this study are available upon reasonable request. MATLAB codes for motion-corrected PALM are available at https://github.com/PuchnerLab/mcPALM, including an interactive GUI for non-expert users. All analyses were run in MATLAB 2024b.

## Acknowledgements

The authors acknowledge the International Institute for Biosensing (IIB) at the University of Minnesota for funding and supporting research related to this paper. The authors thank Dushyant Mehra and Angel Mancebo for their valuable help on motion-correction codes and channel-calibration setup.

## References

Adhikari, S., J. Moscatelli, and E.M. Puchner. 2021. Quantitative live-cell PALM reveals nanoscopic Faa4 redistributions and dynamics on lipid droplets during metabolic transitions of yeast. Mol. Biol. Cell. 32:1565–1578. doi:10.1091/mbc.E20-11-0695.

Adhikari, S., J. Moscatelli, E.M. Smith, C. Banerjee, and E.M. Puchner. 2019. Single-molecule localization microscopy and tracking with red-shifted states of conventional BODIPY conjugates in living cells. Nat. Commun. 10:3400. doi:10.1038/s41467-019-11384-6.

Arlt, H., K. Auffarth, R. Kurre, D. Lisse, J. Piehler, and C. Ungermann. 2015. Spatiotemporal dynamics of membrane remodeling and fusion proteins during endocytic transport. Mol. Biol. Cell. 26:1357–1370. doi:10.1091/mbc.E14-08-1318.

Balderhaar, H.J.K., and C. Ungermann. 2013. CORVET and HOPS tethering complexes **–** coordinators of endosome and lysosome fusion. J. Cell Sci. 126:1307–1316. doi:10.1242/jcs.107805.

Banerjee, C., D. Mehra, D. Song, A. Mancebo, J.-M. Park, D.-H. Kim, and E.M. Puchner. 2023. ULK1 forms distinct oligomeric states and nanoscopic structures during autophagy initiation. Sci. Adv. 9:39. doi: 10.1126/sciadv.adh4094

Banterle, N., K.H. Bui, E.A. Lemke, and M. Beck. 2013. Fourier ring correlation as a resolution criterion for super-resolution microscopy. J. Struct. Biol. 183:363–367. doi:10.1016/j.jsb.2013.05.004.

Belyy, V., N.-H. Tran, and P. Walter. 2020. Quantitative microscopy reveals dynamics and fate of clustered IRE1α. Proc. Natl. Acad. Sci. 117:1533–1542. doi:10.1073/pnas.1915311117.

Betzig, E., G.H. Patterson, R. Sougrat, O.W. Lindwasser, S. Olenych, J.S. Bonifacino, M.W. Davidson, J. Lippincott-Schwartz, and H.F. Hess. 2006. Imaging Intracellular Fluorescent Proteins at Nanometer Resolution. Science. 313:1642–1645. doi:10.1126/science.1127344.

Cabrera, M., M. Nordmann, A. Perz, D. Schmedt, A. Gerondopoulos, F. Barr, J. Piehler, S. Engelbrecht-Vandré, and C. Ungermann. 2014. The Mon1-Ccz1 GEF activates the Rab7 GTPase Ypt7 via a longin fold-Rab interface and association with PI-3-P-positive membranes. J. Cell Sci. jcs.140921. doi:10.1242/jcs.140921.

Calebiro, D., F. Rieken, J. Wagner, T. Sungkaworn, U. Zabel, A. Borzi, E. Cocucci, A. Zürn, and M.J. Lohse. 2013. Single-molecule analysis of fluorescently labeled G-protein–coupled receptors reveals complexes with distinct dynamics and organization. Proc. Natl. Acad. Sci. 110:743–748. doi:10.1073/pnas.1205798110.

Casler, J.C., and B.S. Glick. 2020. A microscopy-based kinetic analysis of yeast vacuolar protein sorting. eLife. 9:e56844. doi:10.7554/eLife.56844.

Chang, F.S., G.-S. Han, G.M. Carman, and K.J. Blumer. 2005. A WASp-binding type II phosphatidylinositol 4-kinase required for actin polymerization-driven endosome motility. J. Cell Biol. 171:133–142. doi:10.1083/jcb.200501086.

Chang, F.S., C.J. Stefan, and K.J. Blumer. 2003. A WASp Homolog Powers Actin Polymerization-Dependent Motility of Endosomes In Vivo. Curr. Biol. 13:455–463. doi:10.1016/S0960-9822(03)00131-3.

Christoforidis, S., M. Miaczynska, K. Ashman, M. Wilm, L. Zhao, S.-C. Yip, M.D. Waterfield, J.M. Backer, and M. Zerial. 1999. Phosphatidylinositol-3-OH kinases are Rab5 effectors. Nat. Cell Biol. 1:249–252. doi:10.1038/12075.

Day, K.J., J.C. Casler, and B.S. Glick. 2018. Budding Yeast Has a Minimal Endomembrane System. Dev. Cell. 44:56–72.e4. doi:10.1016/j.devcel.2017.12.014.

Di Paolo, G., and P. De Camilli. 2006. Phosphoinositides in cell regulation and membrane dynamics. Nature. 443:651–657. doi:10.1038/nature05185.

Gerrard, S.R., N.J. Bryant, and T.H. Stevens. 2000. *VPS21* Controls Entry of Endocytosed and Biosynthetic Proteins into the Yeast Prevacuolar Compartment. Mol. Biol. Cell. 11:613–626. doi:10.1091/mbc.11.2.613.

Gillooly, D.J. 2000. Localization of phosphatidylinositol 3-phosphate in yeast and mammalian cells. EMBO J. 19:4577–4588. doi:10.1093/emboj/19.17.4577.

Girao, H., M.-I. Geli, and F.-Z. Idrissi. 2008. Actin in the endocytic pathway: From yeast to mammals. FEBS Lett. 582:2112–2119. doi:10.1016/j.febslet.2008.04.011.

He, K., R. Marsland Iii, S. Upadhyayula, E. Song, S. Dang, B.R. Capraro, W. Wang, W. Skillern, R. Gaudin, M. Ma, and T. Kirchhausen. 2017. Dynamics of phosphoinositide conversion in clathrin-mediated endocytic traffic. Nature. 552:410–414. doi:10.1038/nature25146.

Hess, S.T., T.P.K. Girirajan, and M.D. Mason. 2006. Ultra-High Resolution Imaging by Fluorescence Photoactivation Localization Microscopy. Biophys. J. 91:4258–4272. doi:10.1529/biophysj.106.091116.

Hirano, M., R. Ando, S. Shimozono, M. Sugiyama, N. Takeda, H. Kurokawa, R. Deguchi, K. Endo, K. Haga, R. Takai-Todaka, S. Inaura, Y. Matsumura, H. Hama, Y. Okada, T. Fujiwara, T. Morimoto, K. Katayama, and A. Miyawaki. 2022. A highly photostable and bright green fluorescent protein. Nat. Biotechnol. 40:1132–1142. doi:10.1038/s41587-022-01278-2.

Horazdovsky, B.F., G.R. Busch, and S.D. Emr. 1994. VPS21 encodes a rab5-like GTP binding protein that is required for the sorting of yeast vacuolar proteins. EMBO J. 13:1297–1309. doi:10.1002/j.1460-2075.1994.tb06382.x.

Huang, B., M. Bates, and X. Zhuang. 2009. Super-Resolution Fluorescence Microscopy. Annu. Rev. Biochem. 78:993–1016. doi:10.1146/annurev.biochem.77.061906.092014.

Huckaba, T.M., A.C. Gay, L.F. Pantalena, H.-C. Yang, and L.A. Pon. 2004. Live cell imaging of the assembly, disassembly, and actin cable–dependent movement of endosomes and actin patches in the budding yeast, *Saccharomyces cerevisiae*. J. Cell Biol. 167:519–530. doi:10.1083/jcb.200404173.

Izeddin, I., V. Récamier, L. Bosanac, I.I. Cissé, L. Boudarene, C. Dugast-Darzacq, F. Proux, O. Bénichou, R. Voituriez, O. Bensaude, M. Dahan, and X. Darzacq. 2014. Single-molecule tracking in live cells reveals distinct target-search strategies of transcription factors in the nucleus. eLife. 3:e02230. doi:10.7554/eLife.02230.

Lachmann, J., C. Ungermann, and S. Engelbrecht-Vandré. 2011. Rab GTPases and tethering in the yeast endocytic pathway. Small GTPases. 2:182–186. doi:10.4161/sgtp.2.3.16701.

Langemeyer, L., F. Fröhlich, and C. Ungermann. 2018. Rab GTPase Function in Endosome and Lysosome Biogenesis. Trends Cell Biol. 28:957–970. doi:10.1016/j.tcb.2018.06.007.

Lee, S.-H., J.Y. Shin, A. Lee, and C. Bustamante. 2012. Counting single photoactivatable fluorescent molecules by photoactivated localization microscopy (PALM). Proc. Natl. Acad. Sci. 109:17436–17441. doi:10.1073/pnas.1215175109.

Mancebo, A., L. DeMars, C.T. Ertsgaard, and E.M. Puchner. 2020. Precisely calibrated and spatially informed illumination for conventional fluorescence and improved PALM imaging applications. Methods Appl. Fluoresc. 8:025004. doi:10.1088/2050-6120/ab716a.

Marat, A.L., and V. Haucke. 2016. Phosphatidylinositol 3-phosphates—at the interface between cell signalling and membrane traffic. EMBO J. 35:561–579. doi:10.15252/embj.201593564.

Markgraf, D.F., F. Ahnert, H. Arlt, M. Mari, K. Peplowska, N. Epp, J. Griffith, F. Reggiori, and C. Ungermann. 2009. The CORVET Subunit Vps8 Cooperates with the Rab5 Homolog Vps21 to Induce Clustering of Late Endosomal Compartments. Mol. Biol. Cell. 20:5276–5289. doi:10.1091/mbc.e09-06-0521.

Mehra, D., S. Adhikari, C. Banerjee, and E.M. Puchner. 2022. Characterizing locus specific chromatin structure and dynamics with correlative conventional and super-resolution imaging in living cells. Nucleic Acids Res. 50:e78–e78. doi:10.1093/nar/gkac314.

Miyashita, M., R. Kashikuma, M. Nagano, J.Y. Toshima, and J. Toshima. 2018. Live-cell imaging of early coat protein dynamics during clathrin-mediated endocytosis. Biochim. Biophys. Acta BBA - Mol. Cell Res. 1865:1566–1578. doi:10.1016/j.bbamcr.2018.07.024.

Mortensen, K.I., L.S. Churchman, J.A. Spudich, and H. Flyvbjerg. 2010. Optimized localization analysis for single-molecule tracking and super-resolution microscopy. Nat. Methods. 7:377–381. doi:10.1038/nmeth.1447.

Mund, M., C. Kaplan, and J. Ries. 2014. Localization microscopy in yeast. In Methods in Cell Biology. Elsevier. 253–271.

Nielsen, E., S. Christoforidis, S. Uttenweiler-Joseph, M. Miaczynska, F. Dewitte, M. Wilm, B. Hoflack, and M. Zerial. 2000. Rabenosyn-5, a Novel Rab5 Effector, Is Complexed with Hvps45 and Recruited to Endosomes through a Fyve Finger Domain. J. Cell Biol. 151:601–612. doi:10.1083/jcb.151.3.601.

Nieuwenhuizen, R.P.J., K.A. Lidke, M. Bates, D.L. Puig, D. Grünwald, S. Stallinga, and B. Rieger. 2013. Measuring image resolution in optical nanoscopy. Nat. Methods. 10:557–562. doi:10.1038/nmeth.2448.

Orr, A., and W. Wickner. 2023. PI3P regulates multiple stages of membrane fusion. Mol. Biol. Cell. 34:ar17. doi:10.1091/mbc.E22-10-0486.

Patki, V., D.C. Lawe, S. Corvera, J.V. Virbasius, and A. Chawla. 1998. A functional PtdIns(3)P-binding motif. Nature. 394:433–434. doi:10.1038/28771.

Peplowska, K., D.F. Markgraf, C.W. Ostrowicz, G. Bange, and C. Ungermann. 2007. The CORVET Tethering Complex Interacts with the Yeast Rab5 Homolog Vps21 and Is Involved in Endo-Lysosomal Biogenesis. Dev. Cell. 12:739–750. doi:10.1016/j.devcel.2007.03.006.

Posor, Y., W. Jang, and V. Haucke. 2022. Phosphoinositides as membrane organizers. Nat. Rev. Mol. Cell Biol. 23:797–816. doi:10.1038/s41580-022-00490-x.

Prescianotto-Baschong, C., and H. Riezman. 1998. Morphology of the Yeast Endocytic Pathway. Mol. Biol. Cell. 9:173–189. doi:10.1091/mbc.9.1.173.

Puchner, E.M., J.M. Walter, R. Kasper, B. Huang, and W.A. Lim. 2013. Counting molecules in single organelles with superresolution microscopy allows tracking of the endosome maturation trajectory. Proc. Natl. Acad. Sci. 110:16015–16020. doi:10.1073/pnas.1309676110.

Rink, J., E. Ghigo, Y. Kalaidzidis, and M. Zerial. 2005. Rab Conversion as a Mechanism of Progression from Early to Late Endosomes. Cell. 122:735–749. doi:10.1016/j.cell.2005.06.043.

Rust, M.J., M. Bates, and X. Zhuang. 2006. Sub-diffraction-limit imaging by stochastic optical reconstruction microscopy (STORM). Nat. Methods. 3:793–796. doi:10.1038/nmeth929.

Seals, D.F., G. Eitzen, N. Margolis, W.T. Wickner, and A. Price. 2000. A Ypt/Rab effector complex containing the Sec1 homolog Vps33p is required for homotypic vacuole fusion. Proc. Natl. Acad. Sci. 97:9402–9407. doi:10.1073/pnas.97.17.9402.

Simonsen, A., R. Lippe, S. Christoforidis, J.-M. Gaullier, A. Brech, J. Callaghan, B.-H. Toh, C. Murphy, M. Zerial, and H. Stenmark. 1998. EEA1 links PI(3)K function to Rab5 regulation of endosome fusion. Nature. 394:494–498. doi:10.1038/28879.

Thompson, R.E., D.R. Larson, and W.W. Webb. 2002. Precise Nanometer Localization Analysis for Individual Fluorescent Probes. Biophys. J. 82:2775–2783. doi:10.1016/S0006-3495(02)75618-X.

Toshima, J.Y., E. Furuya, M. Nagano, C. Kanno, Y. Sakamoto, M. Ebihara, D.E. Siekhaus, and J. Toshima. 2016. Yeast Eps15-like endocytic protein Pan1p regulates the interaction between endocytic vesicles, endosomes and the actin cytoskeleton. eLife. 5:e10276. doi:10.7554/eLife.10276.

Toshima, J.Y., J. Toshima, M. Kaksonen, A.C. Martin, D.S. King, and D.G. Drubin. 2006. Spatial dynamics of receptor-mediated endocytic trafficking in budding yeast revealed by using fluorescent α-factor derivatives. Proc. Natl. Acad. Sci. 103:5793–5798. doi:10.1073/pnas.0601042103.

Tremel, S., Y. Ohashi, D.R. Morado, J. Bertram, O. Perisic, L.T.L. Brandt, M.-K. Von Wrisberg, Z.A. Chen, S.L. Maslen, O. Kovtun, M. Skehel, J. Rappsilber, K. Lang, S. Munro, J.A.G. Briggs, and R.L. Williams. 2021. Structural basis for VPS34 kinase activation by Rab1 and Rab5 on membranes. Nat. Commun. 12:1564. doi:10.1038/s41467-021-21695-2.

Wichmann, H., L. Hengst, and D. Gallwitz. 1992. Endocytosis in yeast: Evidence for the involvement of a small GTP-binding protein (Ypt7p). Cell. 71:1131–1142. doi:10.1016/S0092-8674(05)80062-5.

Zajac, A.L., Y.E. Goldman, E.L.F. Holzbaur, and E.M. Ostap. 2013. Local Cytoskeletal and Organelle Interactions Impact Molecular-Motor-Driven Early Endosomal Trafficking. Curr. Biol. 23:1173–1180. doi:10.1016/j.cub.2013.05.015.

