## Supplementary Materials for "Quantitative Motion-Corrected PALM Links Endosome Structure and Dynamics in Live Cells"

#### 1. Size quantification

In the previous fixed-cell PALM study (Puchner et al., 2013), endocytic vesicles and endosomes were often resolved as donut-like structures. Vesicle sizes could therefore be quantified either from the separation between peaks in mEos2 localization projections or from the standard deviations of mEos2 localization coordinates. In live-cell mcPALM, the effective resolution is lower because motion correction introduces additional uncertainty, as discussed in detail below, making donut-like structures less frequently resolved. We therefore quantified effective endosome size using an effective area enclosing 70% of mEos2 localizations around the localization center of mass (Supplementary Fig. S1B) and validated this approach here.

To account for the angular anisotropy when endosomes are not perfectly circular, localizations were evenly divided into $n$ angular sectors ($n = 20$ in this study). Within each sector, we calculated a radial distance cutoff $d_i$ defined as the distance from the center of mass that enclosed 70% of the localizations in that sector (Supplementary Fig. S1B). The 70% cutoff was chosen as a robust measure of the central localization spread, similar in spirit to standard-deviation-based size estimates. Treating these $d_i$ values as the boundary of an anisotropic area, we defined the effective area $A$ as the sum of all sector areas:

$$A = \sum_{i=1}^{n} \frac{1}{2}\theta {d_i}^2 \qquad (S1)$$

where $\theta = \frac{2\pi}{n}$ is the angular width of each sector. The effective radius was then calculated by equating this effective area to the area of a circle:

$$r = \sqrt{\frac{A}{\pi}} \qquad (S2)$$

This metric reduces the sensitivity to the false-positive localizations near the periphery of the GFP-defined mask while preserving information about the overall localization spread. To validate this size metric, we compared effective radii calculated using this method with radii estimated from standard deviations of mEos2 localization coordinates in both fixed and live cells after motion correction. The two measures were strongly correlated (Supplementary Fig. S1C, linear fit slope = 0.89; Pearson’s r = 0.96), indicating that this effective-radius metric captures the same size trend as the standard-deviation-based approach used in the previous fixed-cell PALM study (Puchner et al., 2013). This supports the robustness of this approach for quantifying endosome sizes in live-cell mcPALM data. The corresponding endosome surface area was then estimated by assuming spherical geometry:

$$A_{surface} = 4\pi r^2 \qquad (S3)$$

#### 2. Size error analysis

##### mEos2 localization uncertainty

Each mEos2 localization was obtained by fitting a single-emitter Point Spread Function (PSF) with a 2D Gaussian function. Its theoretical localization uncertainty was estimated using the Mortensen equation (Mortensen et al., 2010). In the live-cell mcPALM dataset used in Fig. 1, the mean mEos2 localization uncertainty was measured as 23 nm (Supplementary Fig. S2A).

##### GFP centroid determination uncertainty

Because mcPALM corrects mEos2 localizations by subtracting the GFP-derived endosome trajectories, the uncertainty in the GFP centroid determination propagates directly to the corrected mEos2 positions. Estimating this uncertainty is not straightforward because endosomes vary in size and brightness, and their GFP intensity profiles can range from diffraction-limited PSFs to extended non-Gaussian structures.

GFP centroids were therefore determined using two approaches depending on mask size, summed GFP intensity, and whether the GFP intensity profile could be reliably fit as a Gaussian PSF. For small endosomes whose sizes are well below or near the diffraction limit, their GFP signals can be approximated as single-emitter PSFs, and their centroids can be localized by fitting the intensity profile with a 2D Gaussian in Insight3. For these Gaussian-fitted endosomes, we estimated the theoretical centroid uncertainty using the Mortensen equation (Mortensen et al., 2010). However, we still observed a significant fraction of endosomes with sizes comparable to or larger than the diffraction limit and endosomes whose GFP intensity profiles deviated substantially from the 2D Gaussian model (Supplementary Fig. S2B). For these endosomes, including those with GFP-intensity-thresholded mask areas exceeding 49 pixels, corresponding to the 7 × 7 pixel ROI used for single-emitter PSF detection, and those not identified as valid Gaussian PSFs by Insight3, centroids were estimated by calculating the intensity-weighted center of mass over all pixels within the mask.

We then obtained empirical GFP centroid uncertainties across a broad range of endosome mask areas and summed GFP intensities. To do this, we generated calibration data using artificial fluorescent dot arrays projected by a digital micromirror device (DMD) (Mancebo et al., 2020). Under high excitation laser power, these artificial PSFs yielded high signal-to-noise ratios and high localization precision (mean = 5.4 nm, 97% < 7.0 nm), allowing their fitted centers to be treated as ground truth of dot positions (Supplementary Fig. S2C). We then varied the excitation laser power and projected dot size to mimic GFP intensity profiles spanning the observed ranges of endosome mask areas and summed intensities. Additionally, we superimposed experimental background noise recorded from unlabeled yeast strains onto these artificial PSFs. For different combinations of mask areas and intensities, we measured the centroid deviation relative to the ground truth and defined the 70th percentile of the deviation distribution as the representative centroid determination uncertainty. These values allowed us to build a size-intensity-uncertainty map that provides a lookup table for empirically estimating GFP centroid determination uncertainty (Supplementary Fig. S2D). The resulting GFP centroid determination uncertainty distribution assigned to all mEos2 localizations in the live-cell mcPALM dataset used in Fig. 1 yielded a mean of 33 nm with a long tail toward large uncertainty values (Supplementary Fig. S2D).

##### Channel transformation uncertainty

Another source of uncertainty in mcPALM arises from registering the spatially split GFP and mEos2 emission channels with a channel transformation equation group (see Materials and Methods). To quantify the local transformation uncertainty associated with the fitted transformation equations, we moved the DMD-projected dot array to sample different locations across the field of view. For each dot, we transformed its coordinate from the mEos2 channel into the GFP channel using the third-order polynomial transformation equations (Mancebo et al., 2020), and then computed the deviation between the transformed coordinate and the corresponding dot coordinate measured directly in the GFP channel. The deviations from all projected dots were used to generate a spatial map of the local channel transformation uncertainty across the full field of view (Supplementary Fig. S2E). The resulting transformation uncertainty distribution from the live-cell mcPALM dataset used in Fig. 1 had a mean of 36 nm (Supplementary Fig. S2E).

##### GFP centroid interpolation uncertainty

Another source of uncertainty in live-cell mcPALM is introduced by interpolation of the GFP centroid trajectory. To minimize bleed-through between two channels and reduce GFP photobleaching, GFP was excited once every five frames, while mEos2 was excited in the intervening frames. For the 20 Hz movies used in this study, this acquisition scheme corresponds to GFP excitation at 4 Hz, or one GFP frame every 250 ms. Because endosomes continue to move between GFP frames, the GFP centroid position must be estimated for each intermediate mEos2 frame. We used linear interpolation between adjacent GFP frames to assign the GFP centroid position at these intermediate time points. Below, we justify this choice and quantify the uncertainty introduced by this procedure.

Assuming an endosome undergoes one-dimensional Brownian motion (random diffusion), its position $X(t)$ can be mathematically described by a standard Wiener process $W(t)$ and a diffusion coefficient $D$ (Karatzas and Shreve, 1998):

$$X(t) = X(0) + \sqrt{2D}W(t) \qquad (S4)$$

Or equivalently in differential form:

$$dX(t) = \sqrt{2D}dW(t) \qquad (S5)$$

Given the definition of standard Wiener process, the increment, i.e. the displacement from time $t$ to $t + \Delta t$, follows a Gaussian distribution with mean at 0 and variance proportional to elapsed time $\Delta t$:

$$X(t + \Delta t) - X(t) = \frac{1}{\sqrt{4\pi D\Delta t}} e^{-\frac{x^2}{4D\Delta t}} \qquad (S6)$$

which implies the mean-squared displacement (MSD) in 1D:

$$< \triangle X^2 \geq < (X(t + \Delta t) - X))^2 \geq 2D\Delta t \qquad (S7)$$

For interpolation, consider two measured GFP centroid positions at times $t_0 = 0$, $t_1 = T$ and positions $X(0) = X_0$, $X(T) = X_T$. For an intermediate time $t \in (0, T)$, the conditional distribution of $X(t)$ given both fixed endpoints is also called a Brownian bridge (Karatzas and Shreve, 1998).

We next derive the expressions for conditional mean and variance of this intermediate position, providing the expected interpolated position and the associated interpolation uncertainty.

From Eq. S4, the positions $X(t)$ and $X(T)$ can be expressed as:

$$X(t) = X_0 + \sqrt{2D}W(t) \quad (S8)$$

$$X(T) = X_0 + \sqrt{2D}W(T) \quad (S9)$$

Since $X(T)$ is already measured, we can also replace Eq. S9 with known variables:

$$X_T = X_0 + \sqrt{2D}W_T \quad (S10)$$

Given the properties of a standard Wiener process, in which $(W(t), W(T))$ is jointly Gaussian with $\mathbb{E}[W(t)] = 0$, $\mathbb{E}[W(T)] = 0$, $Var(W(t)) = t$, $Var(W(T)) = T$ and $Cov(W(t), W(T)) = t$, the conditional mean and variance are therefore calculated as:

$$\mathbb{E}[W(t)|W(T) = W_T] = \mathbb{E}[W(t)] + \frac{Cov\big(W(t), W(T)\big)}{Var[W(T)]}(W_T - \mathbb{E}[W(t)]) = \frac{t}{T}W_T = \frac{(X_T - X_0)t}{\sqrt{2D}T} \quad (S11)$$

$$Var(W(t)|W(T) = W_T) = Var\big(W(t)\big) - \frac{Cov(W(t), W(T))^2}{Var\big(W(T)\big)} = t - \frac{t^2}{T} = t\left(1 - \frac{t}{T}\right) \quad (S12)$$

Converting back to the conditional mean and variance of $X(t)$ gives:

$$\mathbb{E}[X(t)|X(T) = X_T] = X_0 + \sqrt{2D}\mathbb{E}[W(t)|W(T)] = X_0 + \frac{t}{T}(X_T - X_0) \quad (S13)$$

$$Var(X(t)|X(T) = X_T) = 2DVar[W(t)|W(T)] = 2Dt\left(1 - \frac{t}{T}\right) \quad (S14)$$

Defining $\alpha = \frac{t}{T}(0 < \alpha < 1)$, these expressions become:

$$\mathbb{E}[X(t)|X(T) = X_T] = X_0 + \alpha(X_T - X_0) \quad (S15)$$

$$Var(X(t)|X(T) = X_T) = 2D\alpha(1 - \alpha)T \quad (S16)$$

These results can be extended as single axial component to two-dimensional isotropic Brownian motion with $X(t) = (x(t), y(t))$. The conditional mean remains the same form for each component, while the overall 2D variance becomes:

$$Var(X(t)|X(T) = X_T) = 4D\alpha(1 - \alpha)T \quad (S17)$$

Thus, we find the conditional mean $\mathbb{E}[X(t)|X(T) = X_T]$ is exactly equivalent to the linearly interpolated position between the two endpoints, justifying the use of linear interpolation between the two measured GFP centroids in the mcPALM pipeline. And the conditional standard deviation $\sigma(X(t)|X(T) = X_T)$, i.e. the square root form of conditional variance $\sqrt{Var(X(t)|X(T) = X_T)}$, provides a theoretical prediction of the interpolation uncertainty. Based on Eq. S17, the interpolation uncertainty depends on three factors: (1) the diffusion coefficient $D$, which characterizes how fast the target moves; (2) the time gap $T$ between two measured GFP centroids, which is determined by the sampling rate of GFP excitation; (3) the normalized time $\alpha$ of the interpolated point within that time interval, which corresponds to the timing of the intermittent mEos2 frame being interpolated.

To empirically validate this theoretical prediction and quantify interpolation performance, we tested two types of trajectories: (i) simulated trajectories generated by moving DMD-projected dots to follow pure Brownian motion with different diffusion coefficients, and (ii) experimentally measured endosome trajectories. Both datasets were recorded at 20 Hz for ground truth references, and then downsampled to different time gaps for GFP excitation, including the GFP sampling rate (4 Hz) used in mcPALM. We performed linear interpolation between adjacent downsampled points and computed the deviation between the interpolated positions and the corresponding 20 Hz reference positions, which acted as an empirical measurement of GFP centroid interpolation uncertainty (Supplementary Fig. S2F).

We tested the empirical interpolation uncertainty as a function of the three factors, i.e., the time gap $T$, the normalized intermediate time $\alpha = \frac{t}{T}$ and the diffusion coefficient $D$. Using simulated example trajectories, the 70th-percentile of the squared uncertainty $\sigma^2$, which represents the empirical conditional variance, varied with $\alpha$ in a quadratic, symmetric manner, peaking at $\alpha = 0.5$. This trend was observed for two different frame gaps, 8 and 48 frames, with larger uncertainty being observed for the longer time gap. This behavior is in good agreement with Eq. S17 (Supplementary Fig. S2G). We next extracted the maximal interpolation uncertainty at the midpoint $\alpha = 0.5$. At this midpoint, Eq. S17 predicts:

$$\sigma^2{}_{max} = Var(X(t)|X(T) = X_T)_{\alpha=0.5} = DT \qquad (S18)$$

As expected, the measured midpoint squared uncertainty $\sigma^2{}_{max}$ scaled linearly with $T$, and the slope for each example trajectory was close to its diffusion coefficient $D$ (Supplementary Fig. S2H). Pooling all simulated trajectories (n = 38) with different diffusion coefficients and at different time gaps (ranging from 1 to 100 frames) yielded a fitted slope of 1.04 for $\sigma^2{}_{max}$–$DT$ with a Pearson's correlation coefficient of 0.95, showing the strong agreement between the empirical uncertainty measurement from simulated trajectories and the theoretical prediction (Supplementary Fig. S2I).

Real endosome motion in live cells is more complex, as many endosomes do not undergo ideal Brownian motion and often exhibit time-dependent diffusion coefficients, partial directed transport or confined motion. We therefore classified all experimental endosome trajectories into slow, intermediate, and fast diffusion groups and evaluated the averaged interpolation uncertainty distribution within each group (Supplementary Fig. S2J). The 70th-percentile of empirical variance $\sigma^2$ still followed the same quadratic dependence on $\alpha$ with a maximum at the midpoint $\alpha = 0.5$ (Supplementary Fig. S2K). Across all three motility groups, the averaged midpoint variance $\sigma^2{}_{max}$ vs. $DT$ yielded a linearly fitted slope of 0.99 with a Pearson's correlation coefficient of 0.96 (Supplementary Fig. S2L), indicating that the Brownian bridge prediction in Eq. S17 and Eq. S18 still provides a robust approximation for the interpolation uncertainty in live-cell endosome trajectories. We therefore used this theoretical framework to estimate interpolation uncertainty by inputting three parameters: the diffusion coefficient $D$, the time gap $T$ between two measured GFP centroids in one endosome trajectory, and the normalized intermediate time $\alpha$ of the interpolated mEos2 frame. With this theoretical calculation, GFP centroid interpolation uncertainties estimated from the live-cell mcPALM dataset in Fig. 1 had a mean at 22 nm (Supplementary Fig. S2M). For simplicity, a trajectory-level diffusion coefficient was used in this calculation, although local diffusion coefficients should be used in the future to account for time-dependent diffusivity changes.

Overall, Eq. S17 can be used to calculate the interpolation uncertainty introduced when estimating the GFP centroid position at intermediate mEos2 frames. It predicts that interpolation uncertainty increases with a larger diffusion coefficient, a longer time gap between two GFP frames (i.e. reduced sampling frequency of GFP excitation), and an interpolated time point farther from two measured GFP frames (with the maximum uncertainty occurring near the midpoint between the two GFP frames).

##### Summed uncertainty for motion-corrected mEos2 localizations

Given the four uncertainty sources described above, the total uncertainty for each motion-corrected mEos2 localization was then calculated by propagating all four contributions as a root-sum-square error (Supplementary Fig. S3A):

$$\sigma^2 = {\sigma_{mEos2}}^2 + {\sigma_{GFP}}^2 + {\sigma_{tran}}^2 + {\sigma_{interp}}^2 \qquad (S19)$$

Here, $\sigma_{mEos2}$ is the intrinsic mEos2 localization uncertainty, calculated from the Mortensen equation. $\sigma_{GFP}$ is the GFP centroid determination uncertainty, which was either calculated using the Mortensen equation or obtained from the intensity-area-uncertainty map described above (Supplementary Fig. S2D). $\sigma_{tran}$ is the empirical channel transformation uncertainty estimated from the spatial uncertainty map (Supplementary Fig. S2E). And $\sigma_{interp}$ is the GFP centroid interpolation uncertainty, computed as the square root of Eq. S17. The new lateral localization uncertainty from all mEos2 localizations in the live-cell mcPALM dataset used in Fig. 1 yielded a distribution with a mean of 67 nm (Supplementary Fig. S3B), still providing localization precision below the diffraction limit.

##### Propagated uncertainty for endosome radius and surface area

For each endosome, the uncertainty in effective radius was estimated using a Monte Carlo approach (Supplementary Fig. S3C). Each localization was randomly perturbed by adding a random displacement according to its new localization uncertainty, and the effective radius was recalculated with the new localization coordinates. This procedure was repeated for 2000 iterations, generating a distribution of effective radii for each endosome.

The mean radius from the perturbed distributions was larger than the original radius calculated from the unperturbed localizations (Supplementary Fig. S3C), since localization uncertainty adds spatial spread to the localization cloud. We therefore quantified two effects of localization uncertainty on radius estimation. First, the standard deviation (SD) of the Monte Carlo radius distribution was used as a measure of how sensitive the radius estimate is to localization uncertainty. This Monte Carlo radius SD was positively correlated with the mean localization uncertainty of each endosome and had a mean value of 11 nm across all analyzed live-cell endosomes (Supplementary Fig. S3D). Second, we calculated the radius bias, defined as the difference between the mean Monte Carlo radius and the original unperturbed radius. This bias estimates the expected inflation of the measured endosome size caused by localization uncertainty. The radius bias was also positively correlated with the Monte Carlo radius SD and had a mean value of 32 nm (Supplementary Fig. S3D). The corresponding surface area SD was then obtained by propagating the radius uncertainty $\sigma_r$:

$$\sigma_{surfaceA} = 8\pi r \sigma_r \quad (S20)$$

Similarly, the surface area bias was calculated by replacing $\sigma_r$ with the radius bias. When mapped onto the PI3P-surface-area maturation trajectory, larger endosomes tended to show higher surface-area SD and bias, indicating that uncertainty introduced by mcPALM does contribute to the broadening of the live-cell size distribution (Supplementary Fig. S3E).

We also applied the same analysis to fixed-cell PALM data as a benchmark. Fixed-cell samples showed smaller propagated localization uncertainties, with a mean of 29 nm, compared with 67 nm in the live-cell mcPALM dataset (Supplementary Fig. S3B and S3H). Consistent with this lower localization uncertainty, the fixed-cell PI3P-surface-area trajectory showed a narrower distribution and smaller surface area uncertainty and bias (Supplementary Fig. S3I). These results show that localization uncertainty increases the apparent size spread in live-cell mcPALM, but the uncertainty analysis framework provides an interpretable way to quantify both measurement sensitivity and uncertainty-induced size broadening.

#### 3. Single molecule counting and number error analysis

##### Blink correction

An individual mEos2 fluorophore can transiently enter one or multiple short-lived dark states and later return to the emissive state, generating repeated, nonconsecutive localization events (Annibale et al., 2010, 2011; Lee et al., 2012) and causing molecular overcounting. Because these repeated localizations originate from the same mEos2 molecule, they are expected to occur close together in both space and time. A common strategy is therefore to link localizations within defined spatial and temporal thresholds and merge linked localizations into one molecule by averaging their positions with photon-count weighting (Supplementary Fig. S4A) (Puchner et al., 2013; Lee et al., 2012). This approach, however, requires appropriate selection of the spatial and temporal threshold parameters used for linking localizations. More recently, statistical and machine learning approaches have also been developed for blink correction, including methods with parameter-free inputs (Bohrer et al., 2021; Jensen et al., 2022; Nino et al., 2017; Hummer et al., 2016; Rollins et al., 2015).

In this study, photoactivation was kept relatively low to reduce spatial overlap between simultaneously emitting molecules, allowing threshold-based blink correction to be applied. The spatial and temporal thresholds were determined from spatio-temporal cross-correlation analysis of the localization data (Supplementary Fig. S4B). Specifically, spatio-temporal cross-correlation maps were calculated by enumerating all pairs of localizations and measuring their spatial and temporal separations, $|x_i - x_j|$ and $|t_i - t_j|$. Localization pairs were binned along both axes, and the number of all localization pairs whose separations fell within each radial and temporal interval was accumulated. The resulting pair-count histogram was normalized by the ring area of each radial bin, the width of each temporal bin, the total observation area, the total acquisition duration, and the total number of localizations, yielding a dimensionless spatio-temporal correlation map. Blinking events appear as a short-distance and short-time correlation peak near the origin of this map. We therefore selected the spatial and temporal linking thresholds, $\Delta d$ and $\Delta t$, from the decay of this peak along each dimension, as shown in the x-z and y-z projections in Supplementary Fig. S4B. For fixed cells, we used thresholds of $\Delta d$ = 80 nm and $\Delta t$ = 3 s, similar to values previously used for mEos3.2 (Banerjee et al., 2023). For live cells, we used thresholds of $\Delta d$ = 180 nm and

$\Delta t$ = 3 s. The larger spatial threshold used for live-cell data is introduced by single molecule diffusion. Linked localizations were then merged into a single molecule by photon-count-weighted averaging of their positions. After blink correction, the short-distance and short-time correlation peak close to the origin was substantially reduced, confirming that repeated localizations from blinking events were effectively corrected. The average number of blinking events per molecule was 4 in live-cell data (Supplementary Fig. S4E), and was 3 in fixed-cell data. These values are slightly higher than previously reported blinks per mEos2 molecule, which are typically in the range of ~1–2.6 (Annibale et al., 2011; Lee et al., 2012; Lando et al., 2012; Fricke et al., 2015; Sanchez et al., 2019). This difference likely reflects differences in imaging conditions and potential over-merging caused by high local density of FYVE-mEos2 on endosomes. However, the number of uncorrected localizations and the number of blink-corrected molecules remained strongly correlated across endosomes, indicating that blink correction overall effectively reduced overcounting without disrupting the underlying molecule-number trend (Supplementary Fig. S4E).

**Photoactivation-based correction**

In live-cell imaging, endosomes are trackable only for a limited duration, due to endosomes moving out of focus and photobleaching in the GFP reference channel. This leads to significant undercounting of mEos2 molecules on individual endosomes. To correct for this undercounting, we estimated the actual molecule number from the incomplete molecule count detected in a short tracking window, using an approach based on input energy of the 405-nm photoactivation laser. To rationalize this correction, we start from the basic assumption that photoactivation of mEos2 under 405-nm illumination can be modeled as an inhomogeneous Poisson process, whose activation rate at time $t$ is linearly dependent on the 405-nm laser power at the same time (Lee et al., 2012):

$$\lambda(t) = kI(t) \qquad (S21)$$

where $k$ is the photoactivation constant. Assuming that all activated mEos2 molecules will be subsequently excited and detected, the expected cumulative distribution function (CDF) of mEos2 molecules $f(t)$ is equivalent to the cumulative probability that one mEos2 has been activated by time $t$, or the complementary probability that one mEos2 molecule remains unactivated by time $t$ (Daley and Vere-Jones, 2008):

$$f(t) = \frac{n(t)}{N} = P(T \leq t) = 1 - P(T > t) \qquad (S22)$$

where $n(t)$ is the cumulative number of detected mEos2 molecules, $N$ is the total number of mEos2 molecules, and $T$ is the activation time of one molecule. For a Poisson process, $P(T > t)$ is the same as the probability that zero activation events have occurred, i.e., that the event number $N(t)$ is zero at time $t$:

$$P(T > t) = P(N(t) = 0) = e^{-\int \lambda(t)dt} \qquad (S23)$$

where $\lambda(t)$, as the time-dependent activation rate, can be substituted by Eq. S21. Thus, the cumulative fraction, i.e. CDF, of detected mEos2 molecules $f(t)$ from Eq. S22 can be written as:

$$f(t) = 1 - e^{-\int kI(t)dt} = 1 - e^{-kE(t)} \qquad (S24)$$

where $E(t) = \int I(t)dt$ is the cumulative delivered 405-nm energy. This shows that the CDF of mEos2 molecules $f(t)$ follows an exponential curve that is solely dependent on the input 405-nm energy $E(t)$ and the activation constant $k$. We experimentally validated this relationship across several datasets (Supplementary Fig. S4F), supporting that the normalized activation curve is independent of mEos2 expression level. Since the activation constant $k$ may vary across imaging days due to differences in alignment and illumination conditions, a separate $f(t)$-$E(t)$ calibration curve was generated for each imaging day.

This calibration curve was then used to determine the total molecule number for each endosome from the incompletely detected molecule number. Given the trace start time $t_1$, trace disappearance time $t_2$ and detected mEos2 molecule number $n$, $n$ represents only the molecules expected to be activated over the delivered 405-nm dose within that observation window from $t_1$ to $t_2$. The total molecule number $N$ was therefore estimated as:

$$N = \frac{n}{f} = \frac{n}{f(E(t_2)) - f(E(t_1))} \qquad (S25)$$

where $f(E(t_2))$ and $f(E(t_1))$ are fractions from the calibration curve when the input energy is $E(t_2)$ and $E(t_1)$. To simplify computation, we only analyzed endosomes appearing near the beginning of long movies.

We next quantified the uncertainty associated with this photoactivation-based correction. Given the successfully detected fraction $f$ and detected molecule number $n$, the total molecule number will be $N = \frac{n}{f} = n + U$, where $U$ is the number of undetected molecules. If assuming that $f$ can be certainly determined from the calibration curve, the number of unsuccessful detection, $U$, can be modeled using a negative binomial distribution:

$$U \sim NB(r = n + 1, p = f)$$

with probability density function:

$$P(U = N - n; r = n + 1, p = f) = \frac{N}{U} f^r (1 - f)^k \qquad (S26)$$

The standard deviation of this distribution provides a theoretical estimate of the uncertainty during molecule number correction:

$$SD(N; n + 1, f) = SD(U; n + 1, f) = \frac{\sqrt{(n+1)(1-f)}}{f} \qquad (S27)$$

This expression shows that when the detected molecule number $n$ or detected fraction $f$ is small, the corrected total number becomes less reliable. To validate this prediction, we simulated truncated endosome detection curves with different real molecule numbers and different tracking lengths, and found the empirical uncertainty (calculated as the 70th-precentile deviation from the ground truth) in good agreement with the theoretical prediction from Eq. S27 (Supplementary Fig. S4H). For downstream analysis, we defined an acceptance threshold corresponding to an absolute calibration uncertainty of ≤ 50 molecules. Only endosomes within this threshold, meaning corrected molecule counts with predicted uncertainties below the cutoff of 50 molecules, were retained (Supplementary Fig. S4H).

In practice, the detected fraction $f$ is not strictly fixed and also carries uncertainty. In principle, $f$ could be treated as a probability distribution rather than a single value, for example as a beta distribution that could be incorporated into the negative binomial model. This additional uncertainty mainly reflects movie-to-movie variability when generating the averaged calibration curve and endosome-to-endosome variability of their deviation from the calibration curve. To minimize its influence, we excluded endosomes deviating strongly from the calibration curve, whose interquartile-range-normalized root-mean-squared error (NRMSE) ≥ 1.5.

#### 4. References


Annibale, P., M. Scarselli, A. Kodiyan, and A. Radenovic. 2010. Photoactivatable Fluorescent Protein mEos2 Displays Repeated Photoactivation after a Long-Lived Dark State in the Red Photoconverted Form. *J. Phys. Chem. Lett.* 1:1506–1510. doi:10.1021/jz1003523.

Annibale, P., S. Vanni, M. Scarselli, U. Rothlisberger, and A. Radenovic. 2011. Quantitative Photo Activated Localization Microscopy: Unraveling the Effects of Photoblinking. *PLoS ONE*. 6:e22678. doi:10.1371/journal.pone.0022678.

Banerjee, C., D. Mehra, D. Song, A. Mancebo, J.-M. Park, D.-H. Kim, and E.M. Puchner. 2023. ULK1 forms distinct oligomeric states and nanoscopic structures during autophagy initiation. *Sci. Adv.* 9:39. doi: 10.1126/sciadv.adh4094

Bohrer, C.H., X. Yang, S. Thakur, X. Weng, B. Tenner, R. McQuillen, B. Ross, M. Wooten, X. Chen, J. Zhang, E. Roberts, M. Lakadamyali, and J. Xiao. 2021. A pairwise distance distribution correction (DDC) algorithm to eliminate blinking-caused artifacts in SMLM. *Nat. Methods*. 18:669–677. doi:10.1038/s41592-021-01154-y.

Daley, D.J., and D. Vere-Jones. 2008. An introduction to the theory of point processes. volume 2: General theory and structure / D.J. Daley, D. Vere-Jones. Springer, New York.

Fricke, F., J. Beaudouin, R. Eils, and M. Heilemann. 2015. One, two or three? Probing the stoichiometry of membrane proteins by single-molecule localization microscopy. *Sci. Rep.* 5:14072. doi:10.1038/srep14072.

Hummer, G., F. Fricke, and M. Heilemann. 2016. Model-independent counting of molecules in single-molecule localization microscopy. *Mol. Biol. Cell*. 27:3637–3644. doi:10.1091/mbc.e16-07-0525.

Jensen, L.G., T.Y. Hoh, D.J. Williamson, J. Griffié, D. Sage, P. Rubin-Delanchy, and D.M. Owen. 2022. Correction of multiple-blinking artifacts in photoactivated localization microscopy. *Nat. Methods*. 19:594–602. doi:10.1038/s41592-022-01463-w.

Karatzas, I., and S.E. Shreve. 1998. Brownian Motion and Stochastic Calculus. 113. Springer New York, New York, NY.

Lando, D., U. Endesfelder, H. Berger, L. Subramanian, P.D. Dunne, J. McColl, D. Klenerman, A.M. Carr, M. Sauer, R.C. Allshire, M. Heilemann, and E.D. Laue. 2012. Quantitative single-molecule microscopy reveals that CENP-$A^{Cnp1}$ deposition occurs during G2 in fission yeast. *Open Biol.* 2:120078. doi:10.1098/rsob.120078.

Lee, S.-H., J.Y. Shin, A. Lee, and C. Bustamante. 2012. Counting single photoactivatable fluorescent molecules by photoactivated localization microscopy (PALM). *Proc. Natl. Acad. Sci.* 109:17436–17441. doi:10.1073/pnas.1215175109.

Mancebo, A., L. DeMars, C.T. Ertsgaard, and E.M. Puchner. 2020. Precisely calibrated and spatially informed illumination for conventional fluorescence and improved PALM imaging applications. *Methods Appl. Fluoresc.* 8:025004. doi:10.1088/2050-6120/ab716a.

Mortensen, K.I., L.S. Churchman, J.A. Spudich, and H. Flyvbjerg. 2010. Optimized localization analysis for single-molecule tracking and super-resolution microscopy. *Nat. Methods*. 7:377–381. doi:10.1038/nmeth.1447.

Nino, D., N. Rafiei, Y. Wang, A. Zilman, and J.N. Milstein. 2017. Molecular Counting with Localization Microscopy: A Bayesian Estimate Based on Fluorophore Statistics. *Biophys. J.* 112:1777–1785. doi:10.1016/j.bpj.2017.03.020.

Puchner, E.M., J.M. Walter, R. Kasper, B. Huang, and W.A. Lim. 2013. Counting molecules in single organelles with superresolution microscopy allows tracking of the endosome maturation trajectory. *Proc. Natl. Acad. Sci.* 110:16015–16020. doi:10.1073/pnas.1309676110.

Rollins, G.C., J.Y. Shin, C. Bustamante, and S. Pressé. 2015. Stochastic approach to the molecular counting problem in superresolution microscopy. *Proc. Natl. Acad. Sci.* 112. doi:10.1073/pnas.1408071112.

Sanchez, C.P., C. Karathanasis, R. Sanchez, M. Cyrklaff, J. Jäger, B. Buchholz, U.S. Schwarz, M. Heilemann, and M. Lanzer. 2019. Single-molecule imaging and quantification of the immune-variant adhesin VAR2CSA on knobs of Plasmodium falciparum-infected erythrocytes. *Commun. Biol.* 2:172. doi:10.1038/s42003-019-0429-z.

### 5. Supplementary figures

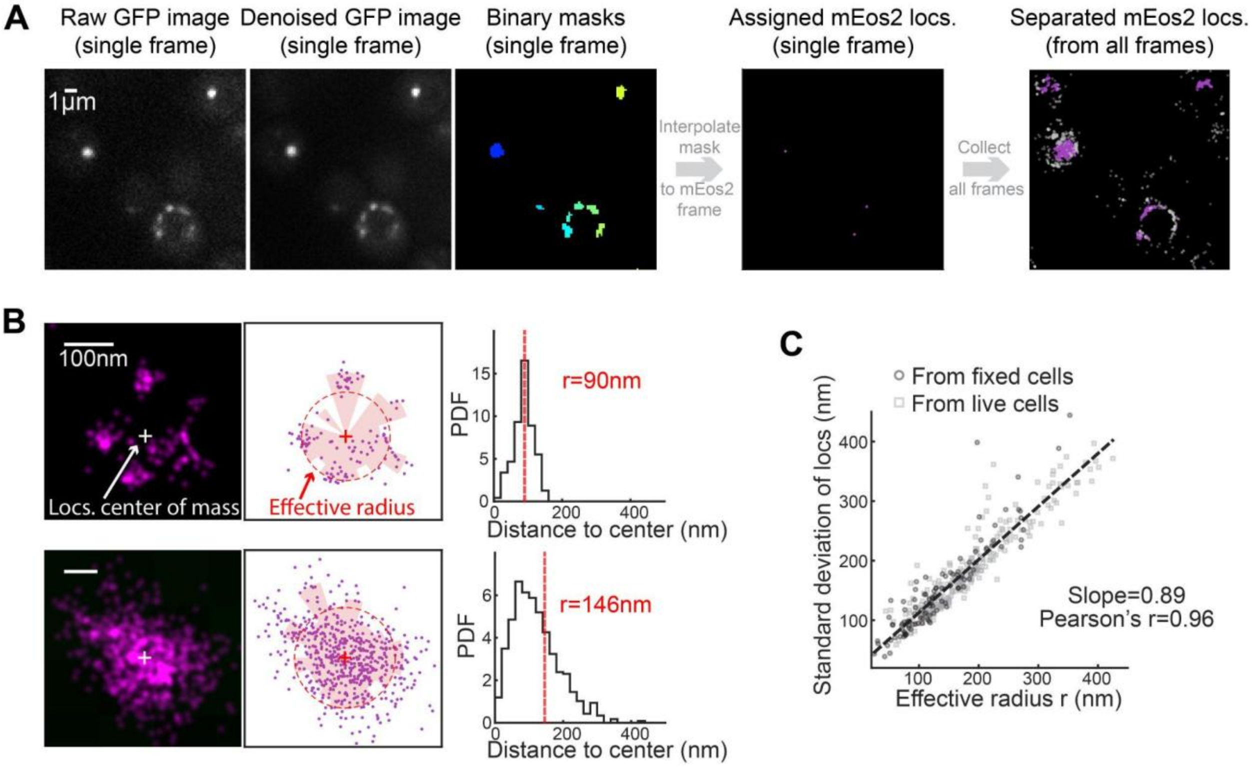


**Supplementary Figure S1. mcPALM image processing and effective radius calculation workflow. (A)** The image segmentation and molecule list splitting pipeline. Raw images from individual GFP frames are denoised using singular value decomposition (SVD) to improve the signal-to-noise ratio. Binary masks are then generated in each image by adaptive thresholding of local GFP intensities to define endosome boundaries for segmentation. The GFP masks are interpolated to the corresponding mEos2 frames, and single mEos2 localizations are assigned to endosomes if they fall within the GFP-defined masks in the corresponding frame. After collecting all frames from one movie, mEos2 localizations assigned to GFP-defined endosomes (shown in magenta) are separated from unassigned localizations that do not colocalize well with detected GFP signals (shown in gray). Approximately 33% of localizations remain unassigned mainly due to the loss of GFP signals when they move out of focus or get photobleached. Scale bar: 1 μm. **(B)** Procedure for calculating the effective radius for representative endosomes from a fixed (top) and a live (bottom) cell. For each endosome, the center of mass ('+') of the PALM localizations (magenta dots) is calculated first. An anisotropic 'effective area' (pink shaded regions) that contains 70% of localizations relative to the center is determined. The effective radius is then calculated as the radius of a circle with the same effective area (red dashed circle). Histograms (right) show the probability density function (PDF) of localization distances to the center and the calculated effective radius (red dashed line). Scale bar: 100 nm. **(C)** Validation of the effective radius as a size metric. The standard deviation of localizations is plotted against the calculated effective radius r for individual endosomes. The strong positive correlation (Pearson's correlation coefficient = 0.96, slope from linear fitting = 0.89) confirms that the effective radius provides a robust estimate of endosome size. Data include 109 endosomes from fixed cells (dark gray, circle) and 228 endosomes from live cells (light gray, square).

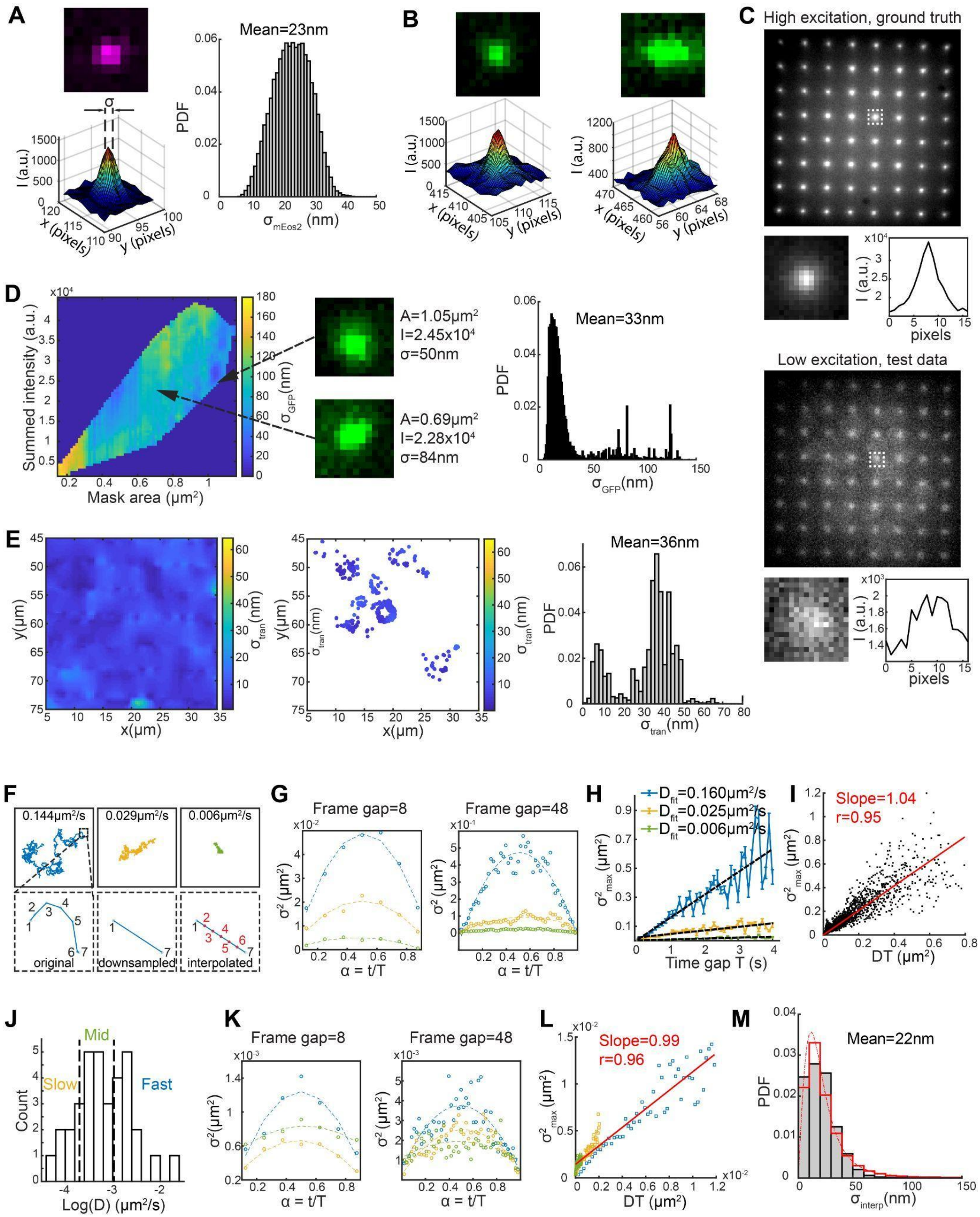


**Supplementary Figure S2. Characterization of four sources of uncertainty: mEos2 localization uncertainty, GFP centroid determination uncertainty, transformation uncertainty, and interpolation uncertainty for mcPALM. (A)** Analysis of FYVE-mEos2 single molecule localization uncertainty $\sigma_{mEos2}$. A magnified view of a single FYVE-mEos2 PSF in the xy-plane (left top) and its corresponding 3D intensity surface plot (left below). $\sigma_{mEos2}$ represents the precision that the PSF center can be determined at. Probability density function (PDF) of $\sigma_{mEos2}$ for all measured localizations in the live-cell mcPALM dataset used in Fig. 1 shows a mean of 23 nm (right). **(B)** Images of two distinct FYVE-GFP spots (top row) from two endosomes with different sizes and intensity profiles are shown along with their respective 3D intensity surface plots (bottom row). Larger or irregular GFP spots can show non-Gaussian intensity profiles, motivating empirical calibration of GFP centroid uncertainty. **(C)** Artificial fluorescent dot arrays used for empirical calibration of the GFP centroid determination uncertainty. An 8 x 8 regular grid pattern is imaged under high excitation to obtain ground-truth dot positions (left top) and under low excitation to generate test data mimicking experimental GFP signal levels (left bottom). Magnified views (right, pixelated images) and their associated 1D intensity profile plots from highlighted regions show the degradation of PSF shape and signal at lower power levels. **(D)** Quantification of empirical GFP centroid determination uncertainty $\sigma_{GFP}$. A 2D scatter plot (left) maps the centroid uncertainty (color-coded) as a function of endosome summed intensity and mask area. Inset images (middle) show examples of two distinct endosomes with their corresponding mask area, intensity, and estimated centroid uncertainty. A histogram (right) shows PDF of $\sigma_{GFP}$ from the live-cell mcPALM dataset used in Fig. 1, yielding a mean of 33 nm. **(E)** Mapping of transformation uncertainty $\sigma_{tran}$ across the full field of view. A 2D heatmap (left) visualizes the spatial distribution of $\sigma_{tran}$ across x and y coordinates of the imaging field of view. A corresponding 2D scatter plot (middle) shows the color-coded $\sigma_{tran}$ of individual localizations at different locations. A histogram (right) shows PDF of $\sigma_{tran}$, with an overall mean of 36 nm. **(F-M)** GFP centroid interpolation uncertainty analysis using simulated (F-I) and experimentally recorded (J-L) high-frame-rate trajectories. **(F)** Simulated trajectories used to evaluate interpolation uncertainty. Top: representative simulated trajectories undergoing Brownian diffusion with different diffusion coefficients, corresponding to fast (blue), intermediate (yellow), and slow (green) motion. Bottom: schematic illustration of interpolation uncertainty arising from downsampling, showing how linear interpolation between sampled positions can deviate from the real trajectory. **(G)** Squared interpolation uncertainty $\sigma^2$, i.e. the variance, calculated from the deviation between the linearly interpolated position and the ground truth position and averaged across individual trajectories, plotted as a function of normalized intermediate time $\alpha = \frac{t}{T}$, for the three example trajectories shown in (F). Results are shown for frame gaps of 8 and 48 between two adjacent GFP frames. $\sigma^2$ follows a symmetric dependence on $\alpha$, with the maximum at the midpoint of the interval. **(H)** Maximum interpolation variance ${\sigma^2}_{max}$ at the midpoint $\alpha = 0.5$ plotted as a function of time gap $T$ for the three example traces in (F). Error bars indicate SEM; dashed lines indicate linear fits. The fitted slopes are close to their diffusion coefficients. **(I)** Relationship between ${\sigma^2}_{max}$ and $DT$ for all simulated trajectories (n = 38) and all sampled time gaps (ranging between 1–100 frames). The relation shows linear scaling with a fitted slope of 1.04 and a Pearson's correlation coefficient of 0.95, consistent with the Brownian-bridge prediction in Eq. S17 and Eq. S18. **(J)** Distribution of diffusion coefficients calculated from 34 experimentally recorded endosome trajectories acquired at 20 Hz. Trajectories were divided into slow, intermediate, and fast diffusion groups, with slow defined as D < $10^{-3.7}$ $\mu m^2/s$, fast defined as D > $10^{-3}$ $\mu m^2/s$, and intermediate defined as values between these thresholds. **(K)** Interpolation variance $\sigma^2$ as a function of $\alpha$ for experimental endosome trajectories, calculated for frame gaps of 8 and 48 and averaged within each diffusion group. **(L)** Relationship between ${\sigma^2}_{max}$ and $DT$ for experimental endosome trajectories, showing an approximately linear dependence with a fitted slope of 0.99 and a Pearson's correlation coefficient of 0.96, again in agreement with the Brownian-bridge prediction in Eq. S17 and Eq. S18. **(M)** Gray: PDF of interpolation uncertainty $\sigma_{interp}$, calculated for all localizations in the live-cell mcPALM dataset used in Fig. 1, using the theoretical equation Eq. S17. The mean interpolation uncertainty is 22 nm. Red: PDF of empirical interpolation uncertainty calculated from 20-Hz reference trajectories in (J), after downsampling with frame gap = 5, matching the sampling rate of mcPALM in this study. The empirical distribution is fitted with a Burr distribution (dotted line) and is shown just as a reference.

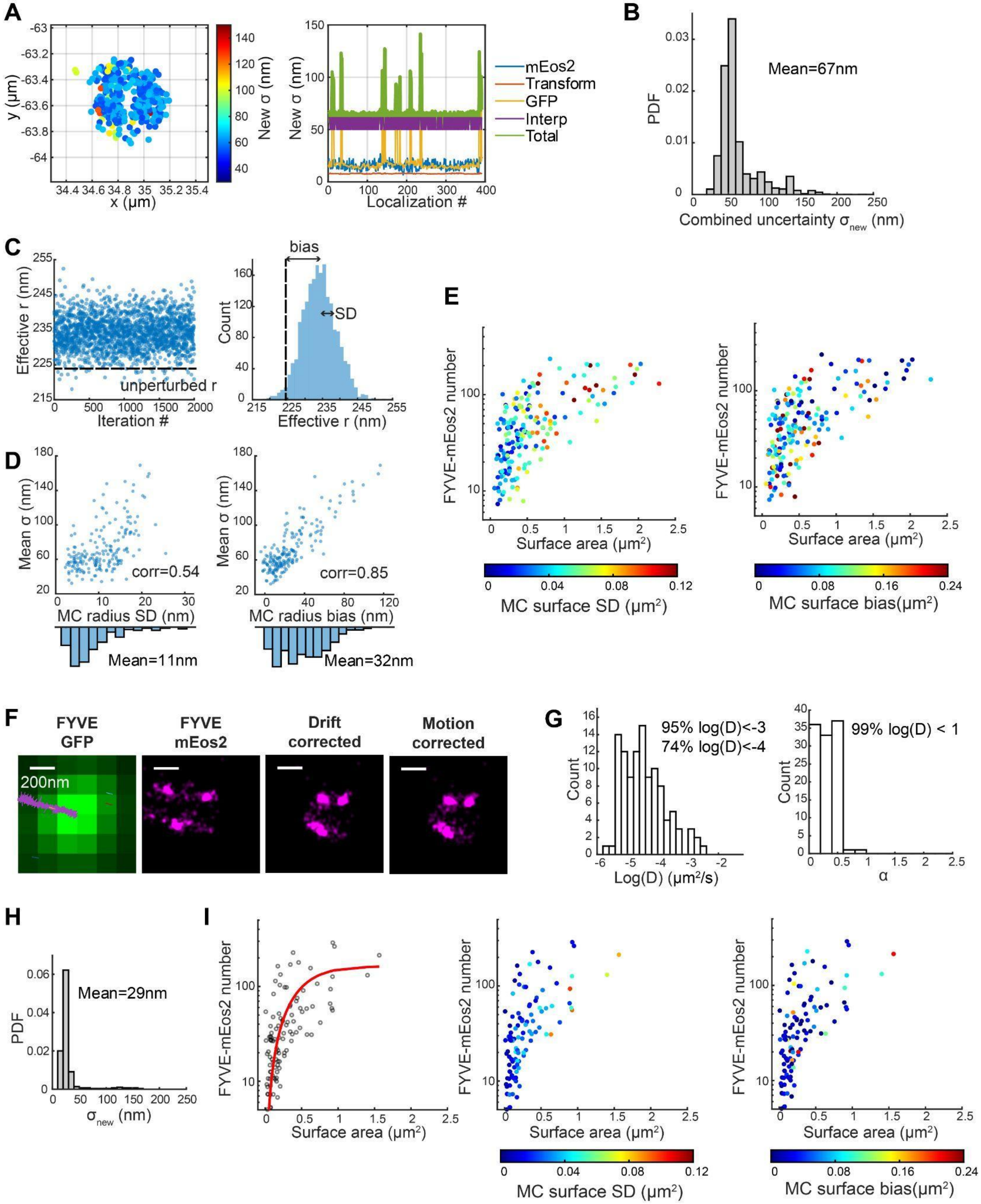


**Supplementary Figure S3. Estimation of new localization uncertainty from the four error sources, Monte Carlo-derived size uncertainty and fixed-cell validation using the same mcPALM analysis pipeline. (A)** FYVE-mEos2 localizations from an example endosome. Left: spatial distribution of localizations with colors indicating the combined localization uncertainty $\sigma_{new}$. Right: decomposition of the localization uncertainty into four sources: mEos2 localization precision (blue), channel transformation uncertainty (orange), GFP centroid determination uncertainty (yellow), and

GFP centroid interpolation uncertainty (magenta). The combined uncertainty is shown in green. **(B)** Probability density function (PDF) of the combined localization uncertainty $\sigma_{new}$ calculated from 58070 FYVE-mEos2 localizations across 193 endosomes. The histogram shows a mean value of 67 nm. **(C)** Monte Carlo estimation of the radius uncertainty for the example endosome shown in (A). Random perturbations were applied to individual localizations according to their combined localization uncertainty, and the effective radius was recalculated for each iteration. Left: effective radius obtained in each Monte Carlo iteration. Right: distribution of effective radii from all iterations. The dashed line indicates the original effective radius measured from the unperturbed localizations. Two effects from localization uncertainties are reported: the radius standard deviation and the radius bias relative to the unperturbed radius. **(D)** Summary of Monte Carlo-derived radius uncertainty across 193 endosomes. Left: relationship between the mean localization uncertainty from all localizations in individual endosomes and the Monte Carlo-derived radius standard deviation shows a positive relation with a Pearson's correlation coefficient = 0.54. Right: relationship between the mean localization uncertainty and the Monte Carlo-derived radius bias with a Pearson's correlation coefficient = 0.85. Histograms show the corresponding distributions, with mean values of 11 nm for the radius standard deviation and 32 nm for the radius bias. **(E)** The same PI3P-surface-area maturation map as in Fig. 2, while color-coded by the surface area standard deviation and bias for each endosome. **(F-I)** Fixed-cell analysis. **(F)** Example of an endosome in a fixed cell but with long stage drift. From left to right: diffraction-limited FYVE-GFP image, raw FYVE-mEos2 PALM image, drift-corrected PALM image, and motion-corrected PALM image. The same motion-correction pipeline used for live-cell analysis is applied here. Scale bar: 200 nm. **(G)** Histograms of the logarithm of the diffusion coefficient, log(D) (left) and anomalous exponent, α (right), from 109 endosome traces recorded in fixed cells. Most traces are effectively fixed and immobile (94.5% with log(D) < -3, 74.1% with log(D) < -4 and 99.1% with α < 1). **(H)** PDF of $\sigma_{new}$ for 37598 FYVE-mEos2 localizations from 109 endosomes. The mean value is 29 nm as indicated in the panel. **(I)** Left: relationship between accessible PI3P content, reported by FYVE-mEos2 molecule number, and endosome surface area, follows the same two-phase maturation sequence observed in live cells but with a narrower size distribution. The red curve shows a guide to the eye. Middle and right: same maturation map as at left, color-coded by the surface area standard deviation and bias. 109 endosomes are shown in the plots.

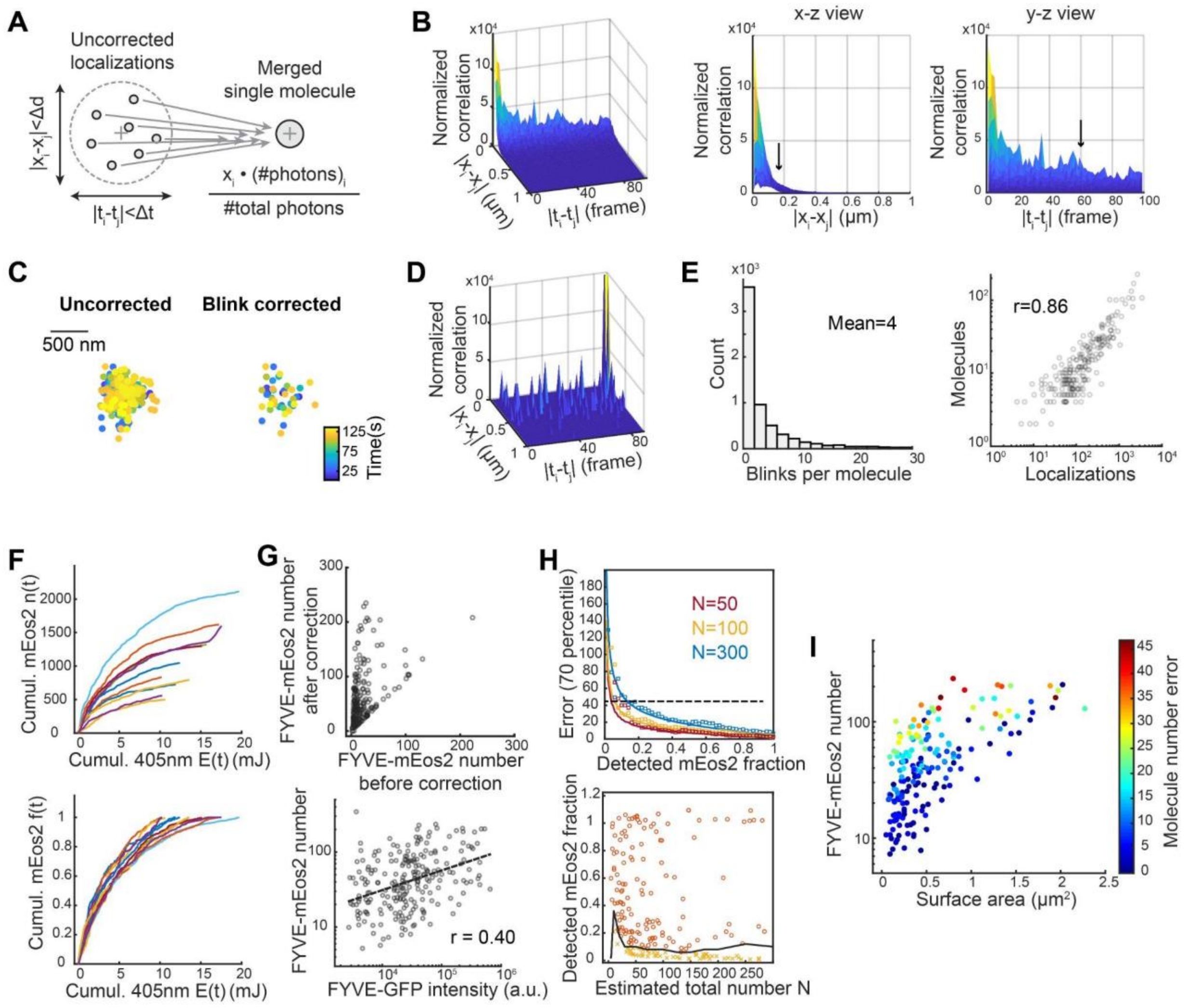


**Supplementary Figure S4. Correction strategies for improving single-molecule counting in live-cell mcPALM.** **(A)** Schematic of the blink correction procedure. Repeated, nonconsecutive localizations arising from the same mEos2 molecule were identified using spatial and temporal thresholds. Linked localizations were merged into a single molecule by photon-count-weighted averaging of their positions. **(B)** Spatial-temporal cross-correlation analysis of uncorrected FYVE-mEos2 localizations in a live-cell movie, showing short-distance and short-time correlations caused by blinking events. The x-z and y-z projections show the correlation along the spatial and temporal axes, respectively. Arrows indicate the selected spatial (180 nm) and temporal thresholds (60 frames or 3 s) used for merging repeated localizations in live cells. **(C)** An example endosome with FYVE-mEos2 localizations before blink correction (left) and merged molecules after blink correction (right). Localizations are color-coded by time. Blink correction reduces repeated localizations from the same emitter while preserving the spatial organization of the endosome. Scale bar: 500 nm. **(D)** Spatial-temporal cross-correlation of blink-corrected molecules from the same list shown in (B), showing reduced short-distance and short-time correlation from blinking events. **(E)** Left: distribution of the number of blinking events per FYVE-mEos2 molecule in 25 live-cell movies, with a mean of 4 calculated from 6266 molecules. Right: correlation between the number of uncorrected FYVE-mEos2 localizations and the number of blink-corrected molecules for 219 endosomes in live cells. Pearson's correlation coefficient = 0.87 indicates a strong correlation. **(F)** Photoactivation-based correction for molecule-number underestimation caused by incomplete tracking windows in live-cell imaging. Top: cumulative photoactivated FYVE-mEos2 number $n(t)$ vs. cumulative 405-nm laser energy $E(t)$. Bottom: cumulative distribution function (CDF) of activated FYVE-mEos2 $f(t)$ vs. cumulative 405-nm laser energy $E(t)$. Curves are generated from 11 movies acquired on the same day. **(G)** Top: comparison of FYVE-mEos2 molecule numbers before and after photoactivation-based correction. Bottom: correlation between corrected FYVE-mEos2 molecule number and FYVE-GFP intensity, with Pearson's correlation coefficient = 0.40. For both plots, each point represents one endosome. A total of 228 endosomes are included. **(H)** Top: 70th-percentile uncertainty from the photoactivation-based correction plotted as a function of the detected mEos2 fraction for three simulated total molecule numbers, N = 50, 100, and 300. Square markers show the simulated data points, while the lines represent the theoretical prediction from the standard deviation of a negative binomial distribution in Eq. S27. The agreement between simulation and theory validates the theoretical prediction from Eq. S27. Bottom: detected FYVE-mEos2 fraction plotted against the estimated total molecule number N. The black curve represents the threshold of an empirical uncertainty = 50 molecules. Orange dots above the threshold show the retained corrected endosomes that have uncertainties ≤ 50 molecules and will be kept for downstream analysis. The yellow dots below the threshold show the failed ones that will be filtered out. **(I)** PI3P-surface-area maturation map color-coded by the estimated molecule number calibration uncertainty for each endosome. Data contain 193 endosomes.

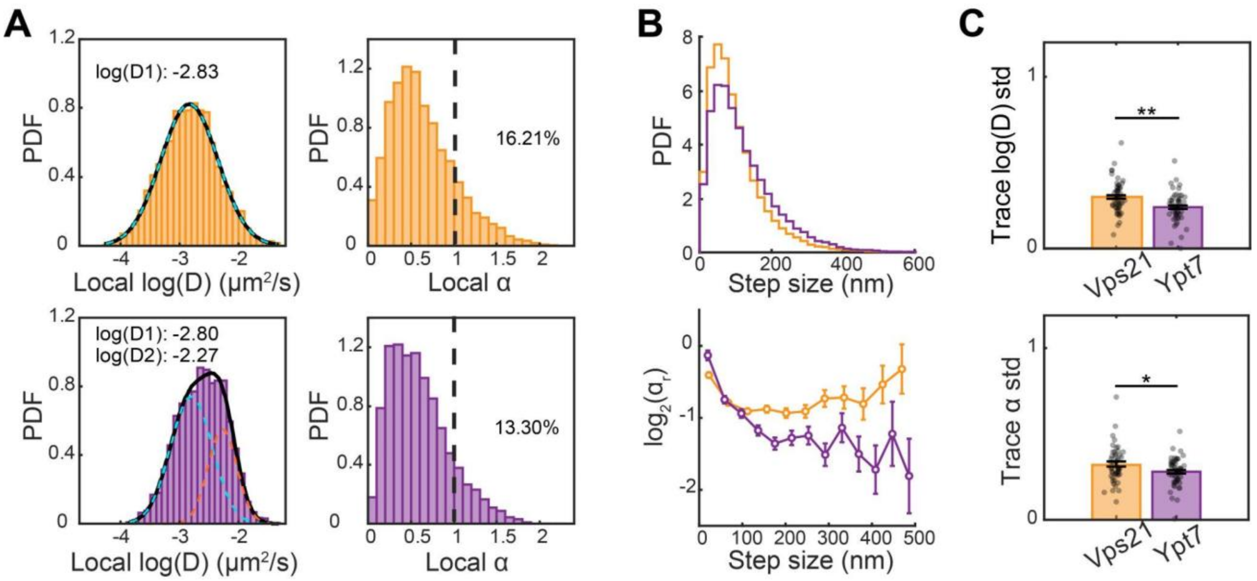


**Supplementary Figure S5. Additional diffusion analysis for Vps21-positive (orange) and Ypt7-positive (magenta) endosomes. (A)** Probability density functions (PDFs) of local diffusivity log(D) (left) and anomalous exponent α (right), calculated using sliding window analysis and segmental MSD fitting as described in Fig. 3 and 4. For fitting log(D) distributions, Gaussian models with one, two or three peaks are selected by Bayesian information criterion (BIC). A single-peak Gaussian with log(D) = -2.83 is chosen for Vps21 data, whereas Ypt7 data is better described by a two-peak Gaussian with log(D) = -2.80 and log(D) = -2.27. The percentages of segments with α > 1 are 16.21% and 13.30% for Vps21 and Ypt7, respectively. The numbers of segments are 23212 and 12643 for Vps21 and Ypt7. **(B)** Top: PDF of step sizes measured over T = 1 s intervals. Step sizes are calculated using Eq. 3. 63601 steps from 50 Vps21-positive endosomes and 26441 steps from 50 Ypt7-positive endosomes are included. Ypt7-positive endosomes show a greater fraction of larger steps than Vps21-positive endosomes. Bottom: the directionality metrics $log_2 a_r$ plotted against step sizes. The step-angle ratio $a_r$ is calculated from Eq. 4. Step sizes are binned, and error bars indicate SEM within each bin. More negative values indicate stronger confinement. Ypt7-positive endosomes show more negative $log_2 a_r$ values at larger step sizes, consistent with stronger long-range confinement. **(C)** Within-trace standard deviations of local log(D) and α. Statistical significance is assessed by one-way ANOVA test, with n.s. no significance, * $p < 0.05$, ** $p < 0.01$, *** $p < 0.001$. Cohen's d values are 0.64 and 0.44, respectively. 50 Vps21-positive endosomes and 50 Ypt7-positive endosomes are included in the statistics.

#### 6. Supplementary videos

**Supplementary Video 1.** Two-color correlative imaging of FYVE-mEos2 (left) in the PALM channel and FYVE-GFP (right) in the conventional channel. Alternating excitation is performed as described in the main text. Scale bar: 5 μm.

**Supplementary Video 2**. Fusion and fission events of FYVE-positive endosomes captured in living yeast cells. Images were averaged over every 10 GFP frames, and the video is played at 10x speed. Scale bar: 1 μm.

**Supplementary Video 3.** Motion of Vps21- and Ypt7-positive endosomes recorded in the GFP channel. The same example endosomes shown in Fig. 5D are used in this video. The corresponding trajectories are color-coded by the time elapsed from the starting point. Images were averaged over every 10 GFP frames just to improve visualization, and the video is played at 10x speed. Scale bar: 1 μm.
